# An *LR* framework incorporating sensitivity analysis to model multiple direct and secondary transfer events on skin surface

**DOI:** 10.1101/2021.02.08.429904

**Authors:** Peter Gill, Øyvind Bleka, Arne Roseth, Ane Elida Fonneløp

## Abstract

Bayesian logistic regression is used to model the probability of DNA recovery following direct and secondary transfer and persistence over a 24 hour period between deposition and sample collection. Sub-source level likelihood ratios provided the raw data for activity-level analysis. Probabilities of secondary transfer are typically low, and there are challenges with small data-sets with low numbers of positive observations. However, the persistence of DNA over time can be modelled by a single logistic regression for both direct and secondary transfer, except that the time since deposition must be compensated by an *offset* value for the latter. This simplifies the analysis. Probabilities are used to inform an activity-level Bayesian Network that takes account of alternative propositions e.g. time of assault and time of social activities. The model is extended in order to take account of multiple contacts between person of interest and ‘ victim’. Variables taken into account include probabilities of direct and secondary transfer, along with background DNA from unknown individuals. The logistic regression analysis is Bayesian -for each analysis, 4000 separate simulations were carried out. Quantile assignments enable calculation of a plausible range of probabilities and sensitivity analysis is used to describe the corresponding variation of *LR*s that occur when modelled by the Bayesian network. It is noted that there is need for consistent experimental design, and analysis, to facilitate inter-laboratory comparisons. Appropriate recommendations are made. The open-source program written in R-code ALTRaP (Activity Level, Transfer, Recovery and Persistence) enables analysis of complex multiple transfer propositions that are commonplace in cases-work e.g. between those who cohabit. A number of case examples are provided. ALTRaP can be used to replicate the results and can easily be modified to incorporate different sets of data and variables.

## 1. Introduction

This paper addresses evaluation of evidence at *activity* level propositions. Experimentation is required in order to inform models with data based upon the transfer, persistence and recovery of DNA. These experiments are intended to simulate events identified by the case circumstances [1], and then the data can be used to derive probabilities that can be used to inform Bayesian Networks (BN).

The specific example of direct skin to skin contact that may occur in an assault is explored here. A previous study by Bowman et al. [2] is compared. The victim may allege that he/she has been held/struck by an assailant and a specific area of skin to skin contact occurred. If the victim and assailant are known to each-other, for example, they may co-habit, then the issue of DNA recovery being affected by social or intimate contacts has to be included in the analysis. DNA will be simultaneously accumulated and lost from surfaces with repeated contact. This process is dynamic and it is important to model the effect.

After the crime-event, the victim reports the crime to the police and samples collected by investigators -swabs will be taken from the affected areas of skin. There will be a time-delay between the crime-event and the collection of skin-swab material. During this time delay, non-self DNA will be lost from the affected area i.e. the longer the time-gap between the crime and the collection of evidence, the less likely it is that DNA will be recovered. The persistence of DNA will be greatly affected if the victim washes the affected area; such activities form an important part of the case circumstances. It can be generalised that after a time period of 24 hours [2] it is much less likely that a DNA profile with high value of sub-source evidence will be recovered from a direct transfer event, but the possibility is not excluded. The aim of this paper is to provide a framework that can model multiple direct and secondary transfer events according to case circumstances based propositions. To achieve this, an existing Bayesian Network (BN) [1, 3] has been adapted. The probabilities to inform the network were derived from experimentation, where DNA recovery, following direct and secondary transfer events were recorded, and the data analysed using logistic regression. The analysis produced continuous probability distributions that were time dependent, thereby taking account of persistence. These logistic regressions measure the decay of DNA recovery over a time period.

A difficulty that is encountered with this kind of study is the limitation of sample sizes and resources required to carry out experiments. Small sample sizes, where DNA recovery of a POI (person of interest) is rarely observed several hours after deposition, are typical with secondary transfer experiments. Fitting probability distributions to data with zero positive observations is problematic. This constraint is not as serious with direct transfer experiments, because sufficient positive observations are achieved provided the time of deposition is less than ≈18hrs (section 6). This facilitates fitting of continuous probability distributions to data, with logistic regression. In order to alleviate the problem of small sample sizes and lack of positive observations with secondary transfer experiments, we have introduced a new method of calculation. This is based upon the principle of two (reasonable) assumptions:

1. The probability that DNA can be recovered from a secondary transfer event is less than that from a direct transfer event.
2. Once DNA has been secondarily transferred to an individual, in the absence of further contact(s), the probability of recovery will decay at a rate that is commensurate with the direct transfer logistic regression curve.

We propose a model for secondary transfer which has the same properties as direct transfer, with the same logistic regression coefficients. However, the time since deposition is re-scaled to take account of the much lower probability of recovery at time zero (section 7).

The model is coded into R and is called Activity Level, Transfer, Recovery and Persistence (ALTRaP) program. It is open-source and freely distributed. It is directly available from https://activitylevel.shinyapps.io/shinyaltrap/. The program opens directly in a browser window; there is no requirement to use or understand the R-environment. Source code (for R-environment), data and user manual are available at: https://sites.google.com/view/altrap/ and https://github.com/peterdgill/altrap. It is hoped that the framework described here is a step towards a standardised method which will facilitate independent analyses by other research groups, to allow comparisons to be made. This will in turn lead to sets of data that can be accessed and utilised by the community to solve complex problems of transfer, persistence and recovery where secondary transfer may be a mechanism. Although the example used here is skin surface, the same framework can be utilised for any example where transfer, persistence and recovery is an issue e.g. on clothing.

It is important to specify that the model presented here can only be used provided that the prosecution sub-source *LR* is accepted by the court. To make clear, a caveat will be needed in the reporting scientist’ s statement. If this is not the case, then evaluation of the evidence at activity level cannot proceed. The model is particularly suited to addressing case circumstances where a ‘ suspect’ and ‘ victim’ are known to each-other, and had ‘ innocent’ social contact some time prior to an assault. For example, they may have met to have a meal, shook hands and embraced. With these kinds of casecircumstances, the mere attribution of DNA to a given individual is not the issue of interest. Rather, the question turns to the probability of its transfer, persistence and recovery after a given time period. To illustrate, two different case circumstances are described and analysed in sections 11 and 12.

## 2. Experimental design

A total of 33 participants were included in the study in different pair combinations. Twelve samples were collected at each time point except 18 and 24 hours, where there was 8 and 6 samples collected from direct and secondary transfer, respectively. This leads to a total of 172 samples in the data-set.

### 2.1. Direct transfer

Participant A was instructed to draw a square of approximately 10 × 5cm on their upper arm. Participant B was then instructed to rub their hand with medium pressure ten times over the marked area of participant A’ s arm, the stroking took place a minimum of one hour after participant B washed hands. The experiment was repeated with sampling of the marked area with a single moistened swab at 0, 1, 3, 6, 9, 12, 18 and 24 hours post-deposit. Only one sample was collected per experiment, since multiple swabbings would adversely affect DNA persistence. A total of 88 samples were collected.

### 2.2. Secondary transfer

A minimum one hour after washing hands, participant A and B shook hands for 30 seconds and then immediately stroked their own upper arm on a pre-marked square 10 × 5cm, ten times with medium pressure. The experiment was repeated with sampling of the marked area with a single moistened swab at 0, 1, 3, 6, 9, 12, 18 and 24 hours post deposit respectively (one sample per experiment). A total of 83 samples were collected

### 2.3. Sample processing

All sampling was performed in clean ‘ DNA-free’ investigation rooms where benches, equipment and hot spots were pre-cleaned with 0.1% hypochlorite solution, and appropriate protection (face mask, hair nets, laboratory coats and gloves) required to enter the rooms. The swabs where removed from the pre-labelled paper bags and the tips were cut by sterile scissors and placed into extraction tubes. The samples were stored at room temperature until DNA was extracted.

DNA extraction was performed using the Chelex^®^ 100 (Bio-Rad Laboratories) protocol (no prior incubation with water) [4], p. 44, using 120*µl* of 5% Chelex solution. All DNA extracts were quantified using DNA Quantifiler™ Trio DNA Quantification kit (Thermo Fisher) with the 7500 Real Time PCR System (Applied Biosystems). PowerPlex^®^ Fusion 6C System (Promega) was used for DNA amplifications, following the manufacturer’ s technical manual for the kit using: 1.0 ng template DNA, 25 mL reaction volume and 29 PCR cycles on Applied Biosystem^®^ Veriti 96-Well Thermal Cyclers (Thermo Fisher). Amplified DNA was subsequently analysed by capillary electrophoresis on the Applied Biosystem^®^ 3500xL Genetic Analyzer (Thermo Fisher), following the PowerPlex^®^ Fusion 6C System Technical Manual (Promega), with an injection time of 24 sec and injection voltage 1.2 kV. Data were analyzed with GeneMapper ID-X v1.2 (Thermo Fisher). The allelic analytical threshold (*AT*) was set to 100 RFU.

### 2.4. Analysis

EuroForMix v.3.1.0 [5] was used to determine sub-source likelihood ratios (*LR*_*ϕ*_), where the numerator was conditioned upon the *POI* and the ‘ victim’ (*V*); the denominator was conditioned upon the ‘ victim’ only. Unknown contributors are designated as *U*_*x*_, where *x* refers to the *x*th unknown. The two alternative propositions are designated *H*_*p*_ as the prosecution proposition and *H*_*d*_ as the defence proposition. *F*_*ST*_ = 0.01 and probability of drop-in = 0.05.

If there are two contributors:

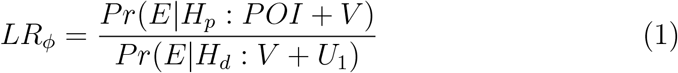

With three contributors:

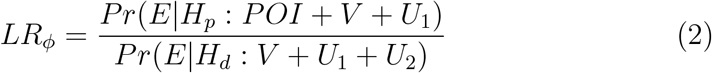

The sub-source likelihood ratio is referred to as *LR*_*ϕ*_ whereas the activity likelihood ratio is *LR*_*a*_; the *H*_*p*_ proposition of *LR*_*ϕ*_ is conditioned as true when activity level is addressed [1].

## 3. Definitions

### 3.1. Transfer, persistence and recovery

A DNA profile that is attributed to a contributor under a given proposition cannot be recovered unless it has been transferred, persisted upon a surface and subsequently recovered in sufficient quantity so that it may be visualised. In accordance with ‘ Consideration 1 and recommendation 8 of [1]: “the scientist can only assign the *probability of recovery* of DNA that is conditioned upon an activity”. In the subsequent text, the term ‘ recovery’ is used in relation to casework to describe the probability that DNA has been transferred, has persisted on an item and has been recovered in sufficient quantities great enough to be genotyped. The term ‘ transfer’ is retained as a general term used to describe the phenomenon or event as it can be observed in experiments where the ground truth is known.

### 3.2. Background DNA

Background DNA present from unknown sources and unknown activities. It can be described as ‘ foreign’ (non-self). We don’ t know how or why it is there [1]. Here background DNA is distinguished from drop-in [6, 7], defined as up to three alleles not observed in prevalent DNA. If four or more alleles are observed, then an ‘ unknown’ contributor is present and *LR*_*ϕ*_ is calculated with eq: (2)

### 3.3. Prevalent DNA

Prevalent DNA is present from known sources/activities that includes ‘ self-DNA’. The analyst has a prior expectation of finding DNA from specific individuals [1].

## 4. List of variables

1. *t* is the probability of direct transfer, persistence and recovery of DNA from the POI (under *H*_*p*_ only).
2. *t*^*′*^ is the probability of direct transfer, persistence and recovery from an unknown assailant (under *H*_*d*_ only).
3. *t*^*′′*^ is the probability of (innocent) direct transfer, persistence and recovery from the POI (where there is common ground agreed under *H*_*p*_ and *H*_*d*_).
4. *b* is the probability of recovering background DNA. Here *b* = 0.36 (under *H*_*p*_ and *H*_*d*_). Based on empirical observation of the data (section 3.2
5. *s* is the probability of secondary transfer, persistence and recovery (under *H*_*p*_ and *H*_*d*_).
6. *h* is the time between deposition of a sample and its collection for analysis. If an assault occurs at 3pm and the samples collected at 10pm then *h* = 7.
7. *n*_*h*_ is the number of contacts modelled within a one hour period of time.
8. *x* is the logistic regression decision threshold based upon *LR*_*ϕ*_.
9. *f*_*x*_ is the offset (time adjustment) applied to the direct logistic regression to convert to a secondary transfer logistic regression, always conditioned on *x*.

### 4.1. Events

Lower case terms, e.g. *t*, are probabilities. Upper case terms, e.g. *T*, signify events, hence *t* is the same as *Pr*(*T*).

## 5. Bayesian Network

The Bayesian Network (BN) described by Taylor et al [3] can be easily adapted to different case circumstances: Gill et al [1], supplement 1, described the same BN in detail, adapted to interpret evidence from underneath fingernails. For the framework described here, the BN is identical to that previously described -the formulae and notation underlying the nodes are the same as described in supplement 1 of [1] except that the probability of recovering background DNA (*b*) is included. The sub-activity nodes are generalised: “X and Y had social contact”, “X assaulted Y” and “Unknown person assaulted Y” (Supplement S1, Fig. S1). The BN will be adapted in later sections to take account of specific case-circumstances. Details of the derivation of formulae and their adaption in the ALTRaP program are provided in the supplementary material S1.

## 6. Direct and secondary transfer data analysis

To inform the BN model outlined above, it is necessary to assign probabilities of DNA persistence and recovery, following secondary and direct transfer. To do this, a series of experimental data (supplement S7, S8) were generated as described in section 2, where the sub-source likelihood ratio (*log*_10_*LR*_*ϕ*_) was compared to the time difference between deposition and sampling (in hours). There is an expectation that longer time difference between two events will result in less DNA that can be recovered from the POI and this will be reflected in reduced probabilities of secondary and direct transfer.

Inspection of scatter-plots of the data (Fig. 1) showed the following:

**Figure 1.**
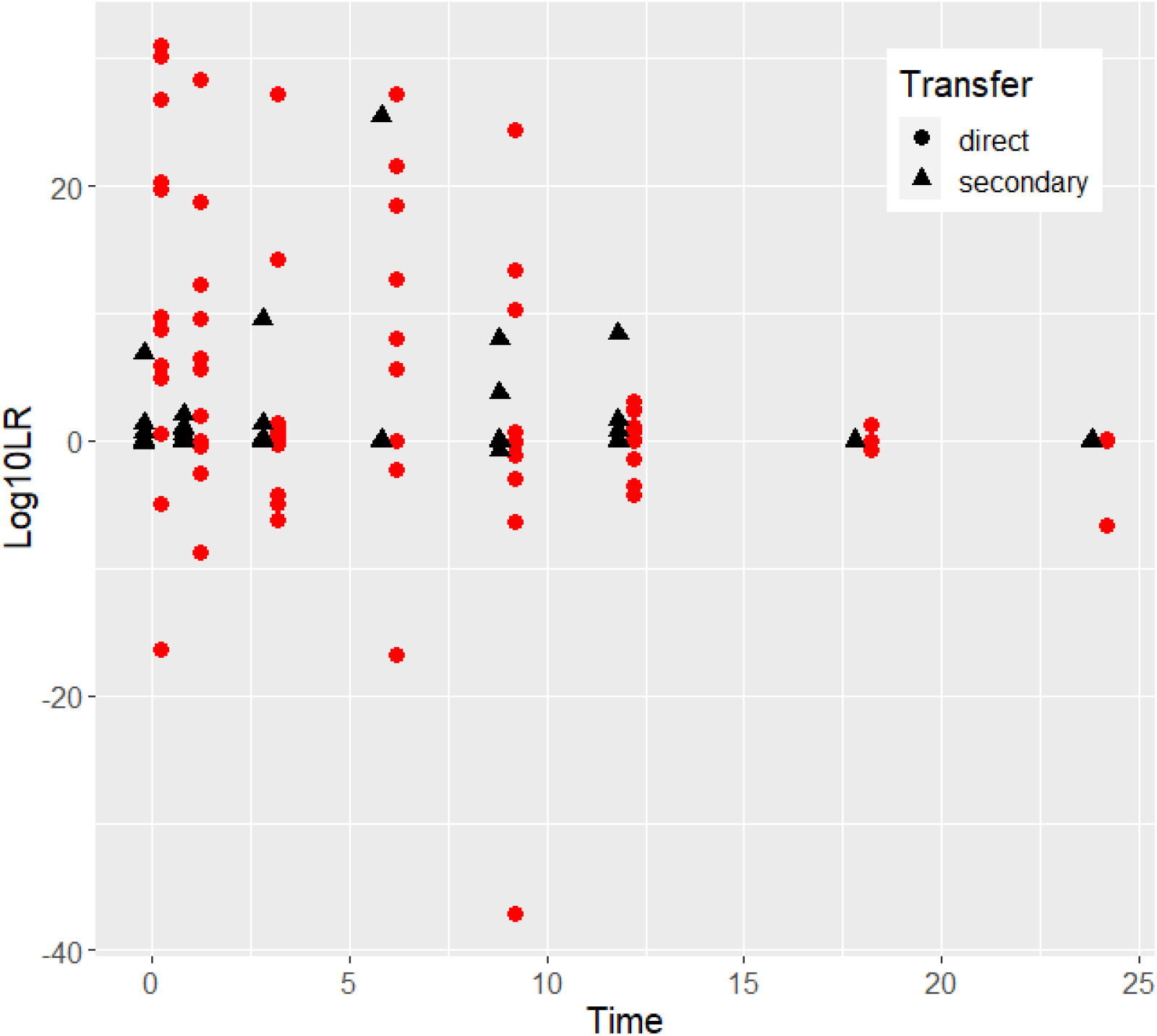
Scatter-plots of direct and secondary transfer showing Time after deposition in hours vs. *log*_10_*LR*_*ϕ*_

### 6.1. Direct transfer

1. There is a wide scatter of points ranging between *log*_10_*LR*_*ϕ*_ *<* 0 to *log*_10_*LR*_*ϕ*_ *≈* 20
2. Observations of higher *LR*_*ϕ*_ values, and their variance diminishes over time. At 24h no *LR*_*ϕ*_ *>* 1 was observed.

### 6.2. Secondary transfer

1. The *LR* values are generally lower than *log*_10_*LR* = 5, but ranged between *log*_10_*LR <* 0 to *log*_10_*LR ≈* 20
2. However, there are a few notable outliers where *LR* values are high, including one at 6 hours where *log*_10_*LR >* 25
3. There were no observations of *LR*_*ϕ*_ *>* 1 when *h ≥* 18

### 6.3. The problem of small data-sets

assigning probabilities from small data-sets are problematic if observations are rare. For the direct transfer data, there was a reasonable spread of observations. For the secondary transfer data, observations of DNA profiles from the POI were much rarer and absent when *h ≥* 18, so that there is a paucity of *LR*_*ϕ*_ *>* 1 observations. This can be rectified by collecting more data, but when probabilities are *Pr* ≈ 0.01 or lower, many samples may be required to achieve an observation of an event and the resource implications become unrealistic to fulfil. Consequently, experimental designs and associated theory are needed to provide a framework that can accommodate the limitations inherent with small data-sets and associated low probabilities. This issue is addressed in section 14.6, where ‘ sensitivity analysis’ is used to examine the outcome from a plausible range of probabilities.

### 6.4. Bayesian logistic regression

To generate probability distributions from the data (supplements S7, S8), logistic regression was used. This method models the probability of a binary event of success vs. no-success: in this case, either a DNA profile is observed from the POI that provides a likelihood ratio ≥*x* or it is *< x*; where *x* is the decision threshold value. From the data, the range of sub-source *LR*_*ϕ*_ s extends from *<* 1 to 10^20^. It is convenient to express *x* in *log*_10_ scale, to be consistent with *log*_10_*LR*_*ϕ*_. The logistic regression coefficients calculate *Pr*(*log*_10_*LR*_*ϕ*_ ≥ *x* |*h*) conditioned upon the time (*h*, in hours) between deposition and sampling.

Using the R stan glm function from package (rstanarm), Bayesian logistic regression utilised MCMC to generate 4000 pairs of coefficients per regression. Package (shinystan) was used in order to check the model diagnostics. A tutorial is provided by [8] and further details provided in Supplement S4.

### 6.5. Direct transfer analysis

Bayesian logistic regression of the data in supplements S7 and S8 was carried out (Fig. 2). Coefficients and goodness of fit tests are shown in Tables 1 and S3. By way of example, at time zero, with *x* = 3 and *x* = 6, respectively: *Pr*(*T* | *h* = 0, *x* = 3) = 0.61 and *Pr*(*T* |*h* = 0, *x* = 6) = 0.52. The persistence and recovery of DNA reduces over time so that at 24 hours the probability assignments are in the region of *Pr* ≈ 0.01.

**Figure 2.**
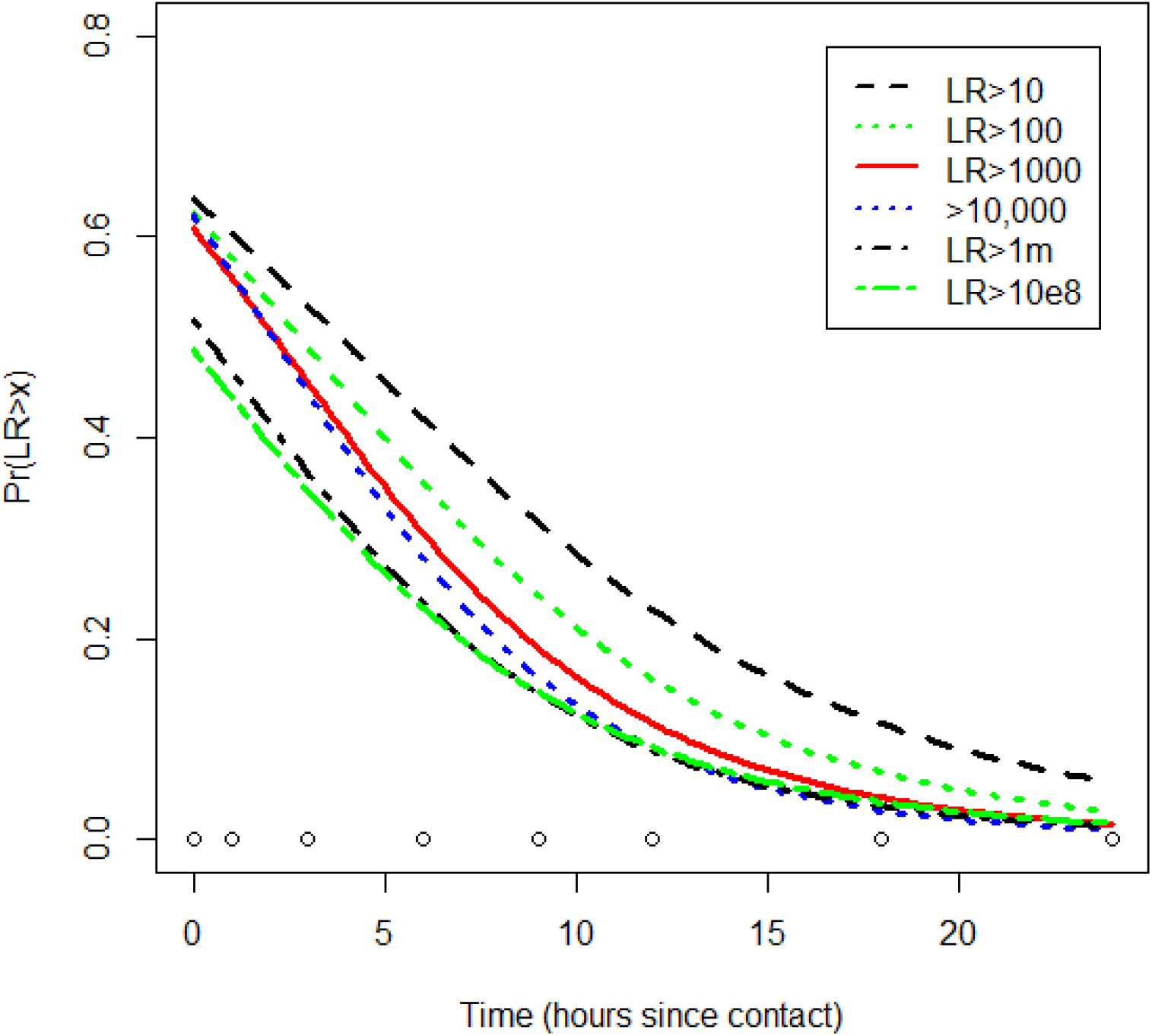
Direct transfer logistic regressions based on median of posterior distribution of coefficients: Time hours since contact and sampling vs. *Pr*(*LR > x*)

**Table 1:**
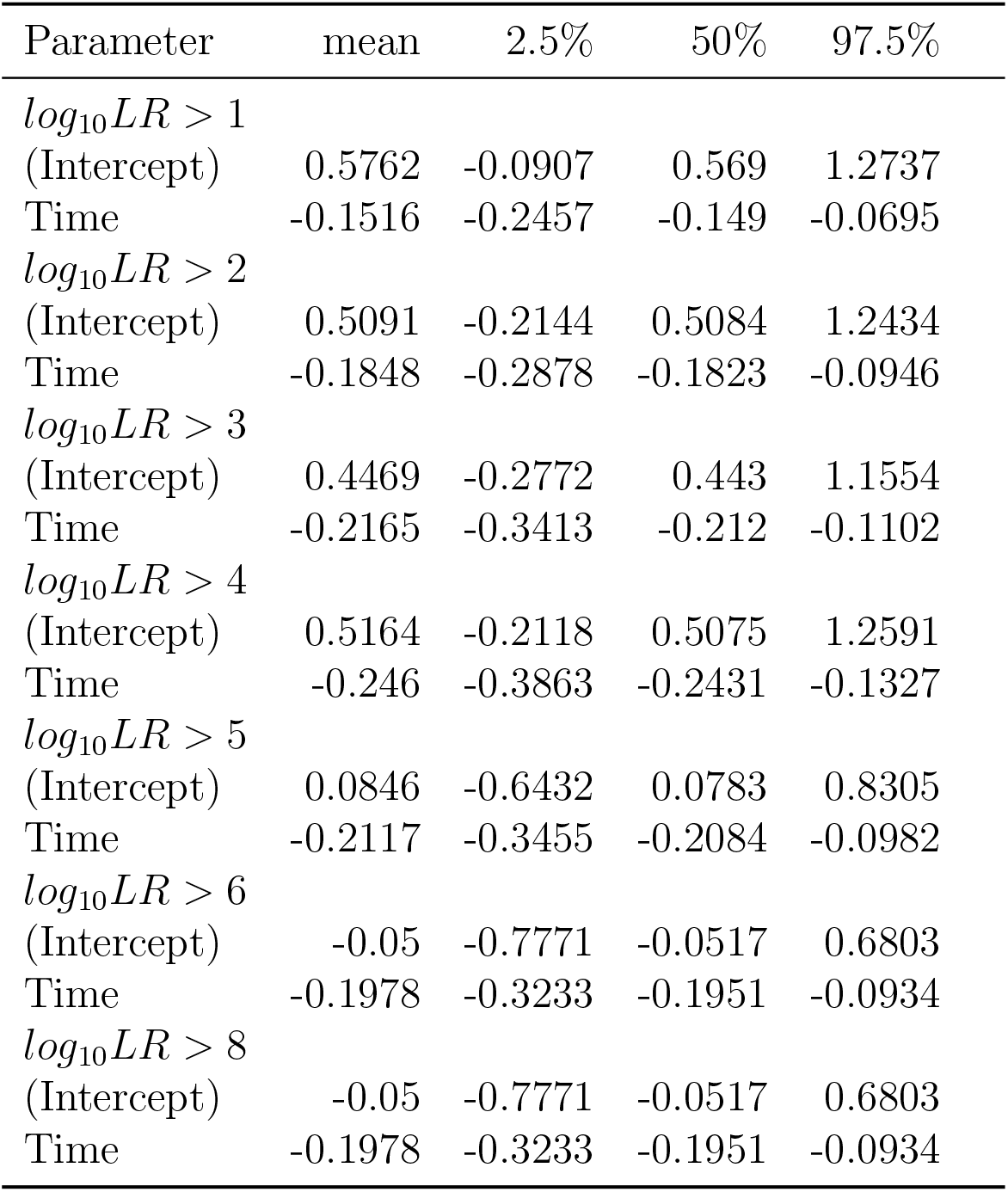
Posterior summary statistics of direct transfer Bayesian logistic regression coefficients generated from 4000 simulations per test showing mean and quantiles. For each decision threshold *x*, there are two coefficients (Intercept and Time) that are used to generate the curves in Fig. 2. An extension of this table is shown in supplement S4.

Summary statistics are shown in Table S3. These tables are posterior summary statistics created using the ‘ shinystan’ application [8]. This application has a host of diagnostic features that can be used to check the fit of the model.

#### 6.5.1. Secondary transfer

Logistic regression was carried out on the secondary transfer data (Fig. S3. However, these lines are almost flat between 0-24 hours; for example the plot for *h* = 1 to *h* = 14 showed that *Pr*(*S* | *x* = 3) dropped from 0.083 -0.048 over the time range. These results violated assumption 1 (section 7), since a *higher* probability of DNA recovery was implied at 24 hours, compared to direct transfer. The model fails, either because there are too few positive data-points to inform the regression curve or because the experimental design was insufficient (see discussion for further comment).

### 6.6. Analysis of background DNA

Defined in section 3.2, from the data of direct and secondary transfer in Supplements S7, S8, the probability of background DNA was calculated to be *b* = 0.36.

## 7. Model assumptions

In the following example, for illustrative purposes, all calculations were made using logistic regressions, where *x* = 3, the logistic regression decision threshold, conditioned on recovery of a probative sub-source DNA profile *log*_10_*LR*_*ϕ*_ *≥* 3.

### 7.1. Assumption 1: For a given time since deposition, Pr(S|h) is always less than Pr(T |h)

Only a small portion of DNA is transferred when a substrate is contacted by a person. Goray et al. [9] showed that 0.36% of biological material is transferred from one site to another, whereas with non-porous substrates approximately 50-95% was transferred. If secondary transfer occurs, in turn, only a small proportion of the DNA from the surface can relocate.

Secondary transfer was measured as probability of recovery of DNA at a specific time(s) since deposition *Pr*(*S* | *h*). For a single contact this is *always* lower than the probability of DNA recovery from a direct transfer event *Pr*(*T* |*h*). From the data, at time *h* = 0, *x* = 3, *Pr*(*S* |*h* = 0, *x* = 3) is approximately 0.05, compared to *Pr*(*T* |*h* = 0, *x* = 3) = 0.61. For any given given time since deposition (*h*) and logistic regression decision threshold (*x*):

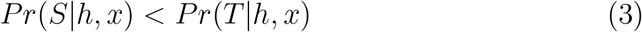

Experimental designs for direct and secondary transfer tests must be com-parable.

### 7.2. Assumption 2: The probability of recovering DNA from secondary transfer follows the same logistic regression as for direct transfer

The working assumption is that for a given threshold value *x*, secondary transfer, *Pr*(*S* |*x*), follows the same logistic regression curve, described by the same pair of coefficients used for direct transfer *Pr*(*T* | *x*). In other words, the decay rate curve for a given *x* value is unaffected by the mode of DNA transfer. This is reasonable because the biological origin of cells propagated by direct or secondary transfer is the same, it is only the much reduced probability of recovery of the latter that needs to be accommodated. Consequently, the decay rate of secondary transfer DNA can be modelled by the same logistic regression curve, but *h* must be adjusted to account for the much lower *Pr*(*S* |*h* = 0, *x*). To carry out the adjustment, an *offset* value is applied (described in next section).

## 8. Assignment of the offset value from secondary transfer data

A set of secondary transfer data (Supplement S8) was generated. The combined data between time zero and one hours after deposition were modelled using a Pareto distribution to determine *Pr*(*S*|*h ∈* 0, 1, *log*_10_*LR*_*ϕ*_ *≥ x*).

Note that *h* = 0 and *h* = 1 were combined so that there were sufficient data to carry out the analysis -afterwards, the offset value was adjusted to *h* = 0. A Pareto model was determined to be the best fit (supplement S3, Fig. S2) to the distribution of *log*_10_*LR*_*ϕ*_ values after testing many different options from R package (fitdistrplus) [10].

### 8.1. Optimising the offset value to calculate Pr(S|h, x)

To obtain *Pr*(*S* | *h, x*) an *offset* value (*f*_*x*_) was calculated. This is a point in time corresponding to the direct transfer logistic regression curve where *Pr*(*T* |*h* = *f*_*x*_, *x*) is the same value as the Pareto distribution assignment of *Pr*(*S* |*h* = 0, *x*). Then the secondary transfer logistic function was calculated from the direct transfer logistic function, using the same coefficients, but time of deposition is adjusted to (*h* + *f*_*x*_) (Fig. 3).

**Figure 3.**
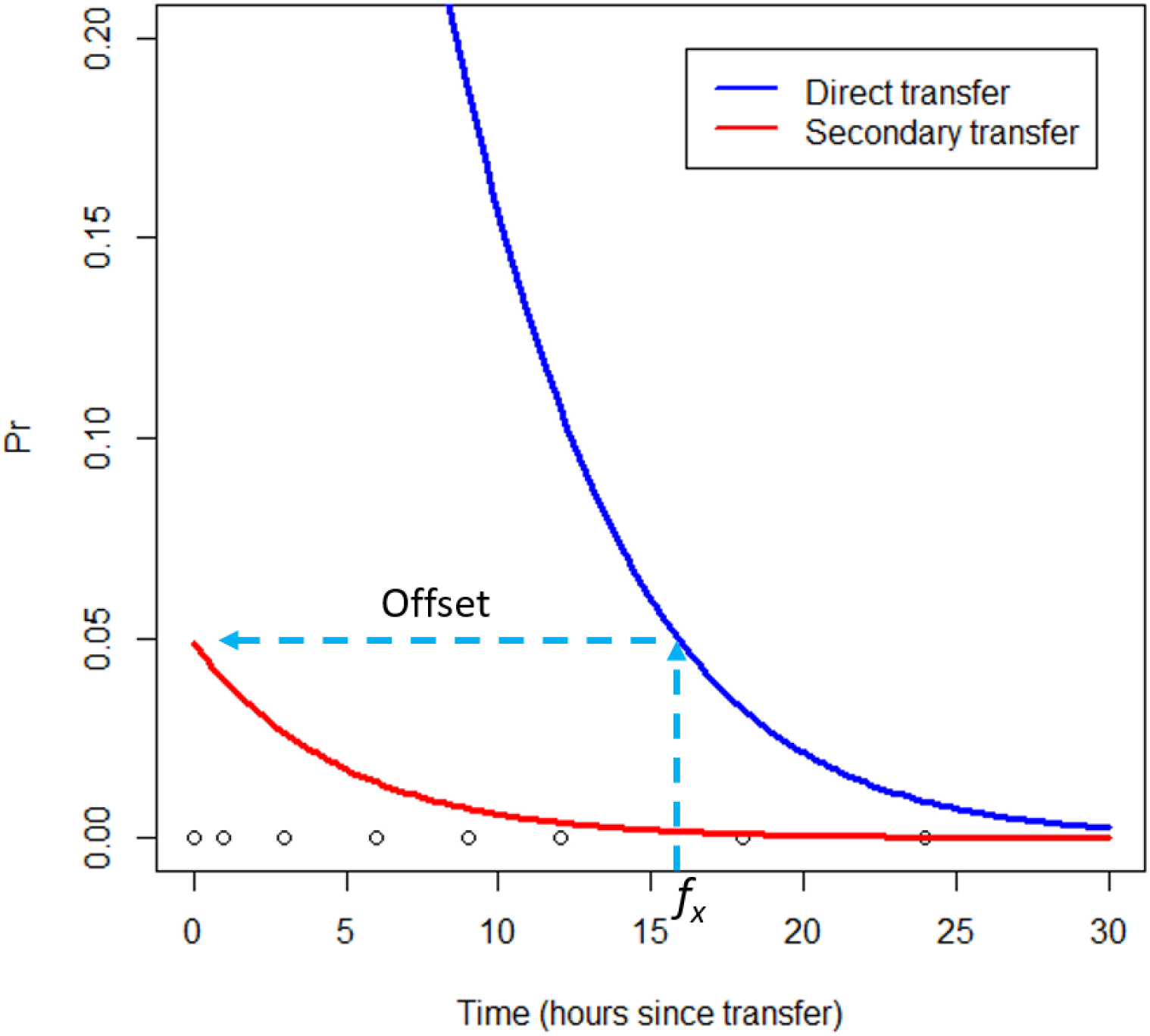
Comparison of probabilities of DNA recovery following direct and secondary transfer vs time between deposition and sampling: *Pr*(*T* | *h*) and *Pr*(*S*|*h*) where *x* = 3. Both direct and secondary transfer logistic regressions employ the same Time and intercept coefficients, but an offset of 15.95 hrs is added to *h* in order to calculate the secondary transfer logistic regression.

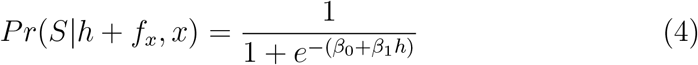

In Fig. 3, the *x*-axis for the secondary transfer logistic function is rescaled so that the secondary transfer derived *h* is the same as direct transfer *h − f*_*x*_ (where *h ≥f*_*x*_). For a given time interval, probabilities for secondary transfer are always less than those for direct transfer and these two types of transfer approach convergence after approximately *h* = 30.

The offset calculation was programmed into ALTRaP as follows:

For a given value of *x*, the corresponding *Pr*(*S* | *h, x*) value was calculated from the Pareto distribution (Fig. S2) of secondary transfer data.

The probability *Pr*(*T* | *h, x*) was calculated using the *β*_0_ and *β*_1_ coefficients, based on posterior medians, from the logistic regression model and the time difference (*h*) between the time of collection of the sample and the time of transfer, for a given choice of *x*. The offset value (*f*_*x*_), in the left hand side of eq. 5 was determined by the exact method described in supplement S5. An alternative method is ‘ optimisation’ using function ‘ optimizer’ from R package (stats). This was used in sensitivity analysis described in section 14.6.

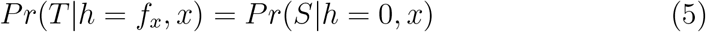

Both *Pr*(*S*) and *Pr*(*T*) are defined by the same *β*_0_ and *β*_1_ logistic regression coefficients calculated from a given *x* value and *Pr*(*S*) is assigned:

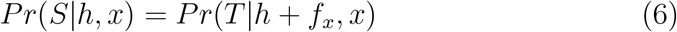

This calculation was carried out for each of the 4000 pairs of logistic regression coefficients generated per sample test, from the posterior distributions of coefficients *β*_0_ and *β*_1_.

For threshold *x* = 3, the mean posterior offset distribution *f*_*x*_*mean*_ = 15.95 hrs. Calculations were repeated for all values of *x* (Table 2). The offset value increased from *f*_*x*_*mean*_ = 15.0 −26.5 hrs as *x* increased from 1 − 10. The secondary and direct transfer decay curves are compared in Fig. 3. Comparison with Fig. S3 shows that the secondary transfer decay curve is no longer flat-lined its entire length, and it is always below the direct transfer decay curve, thereby fulfilling assumption 1 (section 7).

**Table 2:** Secondary transfer, mean (posterior) offset calculation from direct transfer logistic regression.

| $\log_{10}LR > x$ | $f_x$ (hrs) |
| --- | --- |
| 1 | 15.00 |
| 2 | 15.69 |
| 3 | 15.95 |
| 4 | 16.28 |
| 5 | 18.97 |
| 6 | 21.59 |
| 7 | 24.37 |
| 8 | 26.54 |

## 9. The assumption of single transfer events

If an assault is alleged by a victim, a previous innocent social contact, e.g. the victim and suspect attended a party, is often not disputed. This social activity will be included into both *H*_*p*_ and *H*_*d*_ propositions. The difference relates to the assault itself: *H*_*p*_ asserts that the POI assaulted the victim, and there was social activity beforehand, whereas *H*_*d*_ asserts that there was only social activity. To evaluate the evidence, the scientist assigns probabilities of DNA recovery given the time of the assault and the time(s) of social activity, where the latter may include both direct (e.g. a handshake) and secondary transfer (e.g. passing a glass). Under *H*_*d*_ the evaluation of the evidence proceeds under the assumption that the assailant was an unknown individual i.e. no DNA recovery resulting from direct transfer from the POI; recovery of DNA is attributed to secondary transfer only. It is critical to have a good understanding of the times of alleged assault and social activities in order to take into account the effect of DNA loss over time (persistence and recovery). Direct and secondary contacts are not mutually exclusive events -they can occur concurrently. In reality, there may be multiple opportunities at different times for innocent direct and secondary transfer events to occur.

## 10. Modelling multiple transfer events

DNA transfer and accumulation will occur between partners of cohabiting couples,especially where there is intimate contact; promoted by a combination of multiple direct and secondary transfer events [11]. Consequently, it may well be common ground between prosecution and defence that there was opportunity for DNA recovery following direct and/or indirect transfer between a suspect and victim, that was not associated with the crime event. Ideally, a model should accommodate multiple transfer events of different types, at different times. The probability of recovering DNA from skin and/or other surfaces therefore depends upon the frequency and kind of contact with individuals. Regardless of the mode of transfer, over time there are two opposing tendencies: a subject that has continuous or intermittent contact with vectors (a vector is a source) accumulates DNA, whereas no contact with vectors results in loss of DNA.

These counter-opposing forces can be modelled by the binomial expressions described in Supplementary material S2:

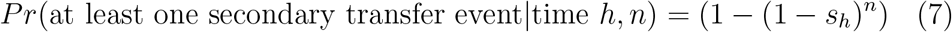

Where *n* is the number of contacts with the vector during a time interval *h*, measured in one hour block (if *h* = 6, this is the start point, meaning that the 1 hour block is between *h* = 6 and *h* = 7 as the end point). In Fig. 4, the time of secondary transfer encompasses four hours and is denoted as *h* = 6, ..10, where *h* = 6 is the end point and *h* = 10 is the end point. To illustrate, a single transfer event can be modelled as: *Pr*(*S* |*h* = 6, *n* = 1) = 0.017. If it was assumed that there were *n* = 4 contacts during a one hour block of time, then *Pr*(*S* |*h* = 6, *n* = 4) = 1*−*(1 −0.017)^4^ = 0.065. The probability *Pr*(*S* |*h, n* = 4) varies according to *h* (see examples in Fig. 4) Consequently, under the assumption of social interaction over a four hour period *h* = 6, .., 10, with four contacts (*n* = 4 per hour); the probabilities per hour block are combined (note *s* = *Pr*(*S*) and *t*^*′′*^ = *Pr*(*T* ^*′′*^):

**Figure 4.**
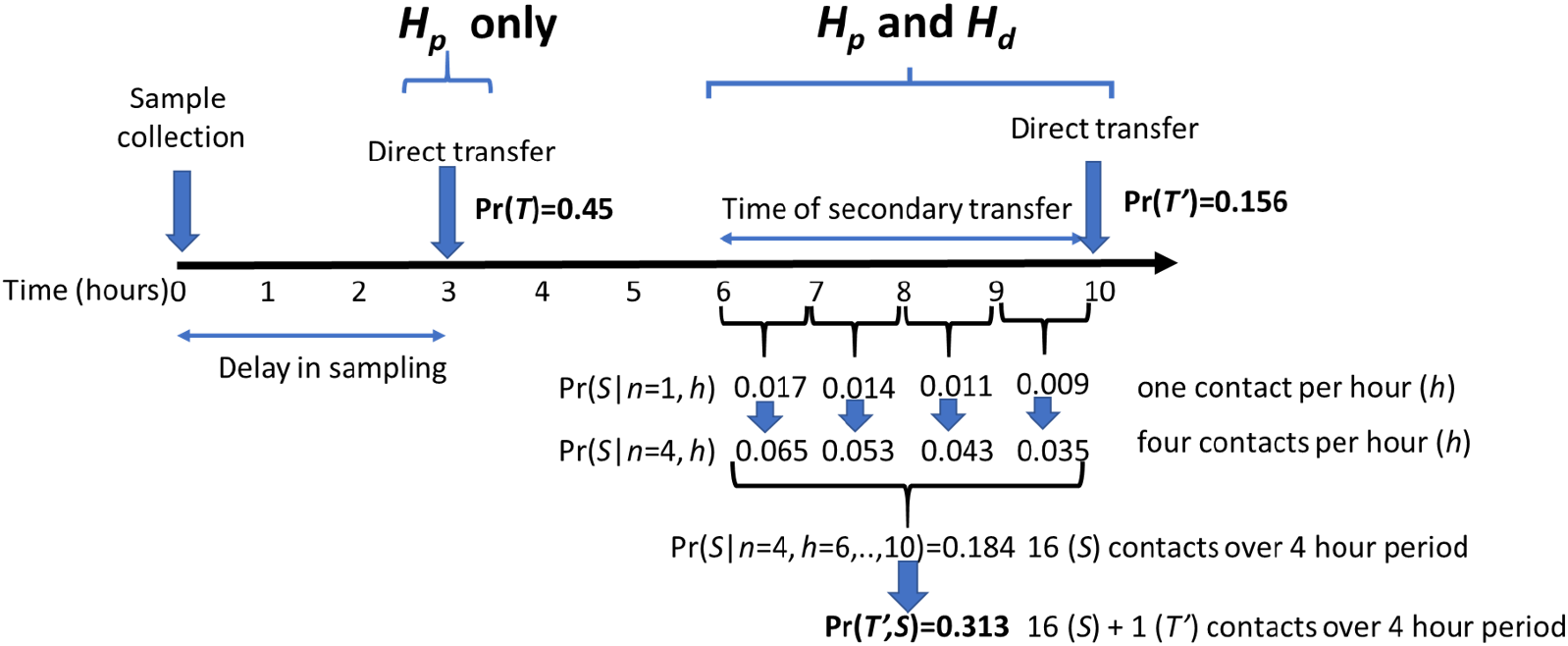
An example showing time since different events occurred, referenced from the time of sampling (*h* = 0), and their conversion into secondary and direct transfer probabilities. There is an assault at time *h* = 3, and social contact at *h* = 6, …, 10, with a direct innocent contact (e.g. handshake) at time 10. There are four contacts per hour, assumed for secondary transfer. Probabilities for each event are shown per hour block, along with their combination.

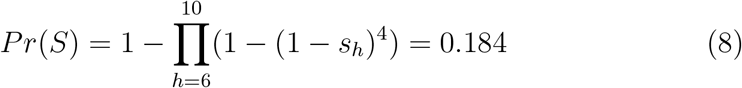

To combine an innocent direct transfer (*T* ^*′′*^) event at time *h* = 10 where *Pr*(*T* ^*′′*^ |*h* = 10, *n* = 1) = 0.156, a binomial expression is utilised which assumes independence between *S* and *T* given *h*:

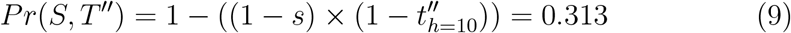

The theory is outlined in Fig. 4 as part of an example that is expanded in the next section. This value is forwarded to the BN calculation described in supplement S2. Note that in the ALTRaP program, calculations are carried out using R package (poibin).

## 11. A worked example

### 11.1. Case circumstances

Ms. X visited Mr. Y at his home, arriving at 10am. They held hands and embraced (this was the only direct contact made) and for a period of 4 hours they socialised, had several drinks, a meal and used the bathroom. After a heated argument she left the house at 2pm and made her way home, a short distance away, arriving at 2.30pm. Leaving her home again at 4pm she decided to go for a walk in the local park where at 5pm she was attacked by a man who approached her from behind, grabbing her arms and knocking her to the ground. He ran away before she could positively identify the attacker. The police were called and samples were taken at 8pm. Mr. Y was apprehended as the prime suspect.

### 11.2. Background information

The victim had showered in the morning at 8am, but did not wash hands or shower before the samples were collected. Before Mr. Y was apprehended, he had fully washed and showered, however.

### 11.3. Analysis

A diagram of events, along with their probabilities are illustrated in Fig. 4. All events are referenced relative to the time that samples were collected, *h* = 0. The assault was at *h* = 3. It was not disputed that the victim had visited the house of the suspect and had social contact with him.

A swab was taken from the victim’ s arm where violent contact had occurred and a DNA profile was obtained. A sub-source likelihood ratio of *LR*_*ϕ*_ = 5000 was calculated with alternative propositions described by eq. 1. However, the sub-source *LR*_*ϕ*_ has no impact upon the activity level propositions which are:

1. Mr. X and Ms. Y had social interaction and she was assaulted by him at the times specified
2. Mr. X and Ms Y only had social interaction at the times specified

Note that to consider activity level propositions, the court would first need to accept that Mr. Y was a donor to the DNA sample which was recovered i.e. the prosecution proposition at sub-source is accepted as being true. The sub-source *LR*_*ϕ*_ can then be used to inform the value of the evidence at activity level via transfer probabilities. The principle of analysing evidence at sub-source and activity levels as two separate consecutive steps is in accordance with ISFG DNA commission recommendations [1].

### 11.4. Results of the analysis

Activity level likelihood ratios (*LR*_*a*_) were calculated using equations eqs. (S6) and (S7) in supplement S1. To model the sensitivity of *LR*_*a*_ to various parameters, different logistic regression threshold decision values of *x* were compared with the effect of *n* = 1*−*4 contacts per hour (Table 3). The highest *LR*_*a*_ was observed when the ‘ POI only’ proposition was tested and undisputed direct transfer at *h* = 10 reduced the value of the *LR*_*a*_. There was also a general increase in *LR*_*a*_ as the logistic regression threshold decision value (*x*) increased.

**Table 3:** Sensitivity analysis of likelihood ratios (*LR*_*a*_) comparing different *log*_10_*LR*_*ϕ*_ *> x* thresholds and between *n* = 1 to 4 secondary transfer events per hour (Fig. 4). Left hand data-set considers effect of undisputed (under *H*_*p*_ and *H*_*d*_) secondary transfer (*S* = 6, …, 10) only. The right-hand data compare *LR*_*a*_s where undisputed secondary (*S* = 6, …, 10) and direct transfer (*T* ^*′′*^ = 10) both occurred. *H*_*p*_ and *H*_*d*_ propositions tested were ‘ POI only’ and ‘ POI and unknown(s)’. The *LR*_*a*_ is always higher for the former.The worked example is shown in bold type.

| | Type of transfer | Secondary ($S$) | | | | Secondary and direct ($S, T''$) | | | |
| --- | --- | --- | --- | --- | --- | --- | --- | --- | --- |
| $Pr(E\vert H_p)$ proposition | No. of contacts per hour<br>$\log_{10} LR > x$ | 1 | 2 | 3 | 4 | 1 | 2 | 3 | 4 |
| POI only | 1 | 5.7 | 3.7 | 3.0 | 2.7 | 3.4 | 2.9 | 2.7 | 2.5 |
|  | 2 | 10.2 | 5.9 | 4.4 | 3.7 | 4.3 | 3.6 | 3.2 | 3.0 |
|  | **3** | **17.8** | **9.6** | **6.9** | **5.5** | **5.2** | **4.5** | **4.0** | **3.7** |
|  | 4 | 33.5 | 17.4 | 12.1 | 9.4 | 6.4 | 5.7 | 5.2 | 4.8 |
|  | 6 | 43.4 | 22.3 | 15.3 | 11.8 | 5.5 | 5.1 | 4.8 | 4.5 |
|  | 8 | 72.5 | 36.9 | 25.0 | 19.1 | 5.2 | 5.0 | 4.8 | 4.7 |
| POI and unknown(s) | 1 | 1.4 | 0.9 | 0.7 | 0.7 | 0.8 | 0.7 | 0.6 | 0.6 |
|  | 2 | 2.8 | 1.6 | 1.2 | 1.0 | 1.2 | 1.0 | 0.9 | 0.8 |
|  | **3** | **5.3** | **2.9** | **2.0** | **1.7** | **1.5** | **1.3** | **1.2** | **1.1** |
|  | 4 | 10.2 | 5.3 | 3.7 | 2.9 | 1.9 | 1.7 | 1.6 | 1.5 |
|  | 6 | 16.5 | 8.5 | 5.8 | 4.5 | 2.1 | 1.9 | 1.8 | 1.7 |
|  | 8 | 29.1 | 14.8 | 10.0 | 7.6 | 2.1 | 2.0 | 1.9 | 1.9 |

The number of secondary transfer contacts per hour was modelled; because this parameter cannot be definitively known, it must be informed by reasonable assumptions about its value as a prior ‘ belief’ based upon the case circumstances. However, the impact was relatively minor. An assessment of *S, T* ^*′′*^ in Table 3 showed the activity *LR*_*a*_ varied between 5.2 *—* 3.7 and 1.5 *−*1.1 for ‘ POI only’ and ‘ POI and unknown(s)’ propositions respectively (*x* = 3). The likelihood ratio for ‘ secondary transfer only’ was always higher than for ‘ secondary and direct’; and it was always higher for ‘ POI only’ compared to ‘ POI and unknown(s), where background DNA was present. The effect of different levels of background DNA on *LR*_*a*_ is described by sensitivity analysis in section 14.7

Such *LR*_*a*_s would be considered as weak support in favour of *H*_*p*_ using the verbal scale described by the ENFSI guideline for evaluative reporting in forensic science [12]. Conversely, if the sub-source logistic regression decision threshold *x* = 6, with one (*S*) contact per hour (‘ POI only’) was assumed with no direct transfer 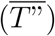 then the *LR* = 43.38 which is moderate support in favour of *H*_*p*_.

## 12. Second example

In real case examples, it is unlikely that the alleged times of direct and secondary events will be the same. For instance, suppose that a victim is assaulted at a party where her arm was violently grabbed, the suspect denies the assault, but claims to have handed a glass of wine to the victim two hours before the alleged assault. There was no other interaction. There was a six hour delay before the victim complained to police and samples were taken from her skin. According to the case circumstances, the social activity that may have led to secondary transfer occurred 8 hours previously. A swab was taken from the victim’ s arm where violent contact had occurred, and a DNA profile was obtained. A sub-source likelihood ratio of *LR*_*ϕ*_ > 1000 was calculated with alternative propositions described by eq. 1. The activity level propositions are the same as for the first example, except that the timings are different:

1. Mr. X and Ms. Y had social interaction and she was assaulted by him at the times specified
2. Mr. X and Ms Y only had social interaction at the times specified

Assuming that the sub-source evidence was accepted by a court, the challenge is to determine the value of the evidence given alternative activity level propositions.

Probability plots of *Pr*(*S*) and *Pr*(*T*) relative to time *h* are illustrated in Fig. 3. Under the assumption that the DNA profile is a mixture of the suspect and the victim, conditioned on six hours for *T*_*h*=6_ and eight hours for *S*_*h*=8_ (*n* = 1), this provided a likelihood ratio of *LR*_*a*_ = 40.8 and *LR*_*a*_ = 18.1 (Table 4), respectively, dependent upon whether the POI was observed with or without unknown contributors. Various other time differences were explored in Table 4 to compare the effect upon *LR*_*a*_. The more distant the time of social contact from the assault the greater *LR*_*a*_ became. This is caused by the decreased probability of *Pr*(*S* |*h*), reflecting the natural loss of foreign DNA from the skin surface over time. It is interesting to note that for ‘ POI only’, the *LR*_*a*_s were similar when the absolute time difference between assault and social interaction was the same: e.g. comparing six and 18 hours with twelve and 24 hours (assault vs social interaction respectively), the *LR*_*a*_ ≈300. However, the probability of actually obtaining a result if *H*_*p*_ is true, approximated by *Pr*(*T*) in Table 4, diminishes from 0.3 (six hours) to 0.11 (twelve hours) i.e. it is important to note that the *LR*_*a*_ provides no information about the probability of obtaining a result, it can only address the probability of the evidence, *if* a result is obtained. For ‘ POI + unknown(s)’, the *LR*_*a*_ is always less than the corresponding ‘ POI only’; the values tend to increase as the absolute time difference between assault and social interaction increases.

**Table 4:**
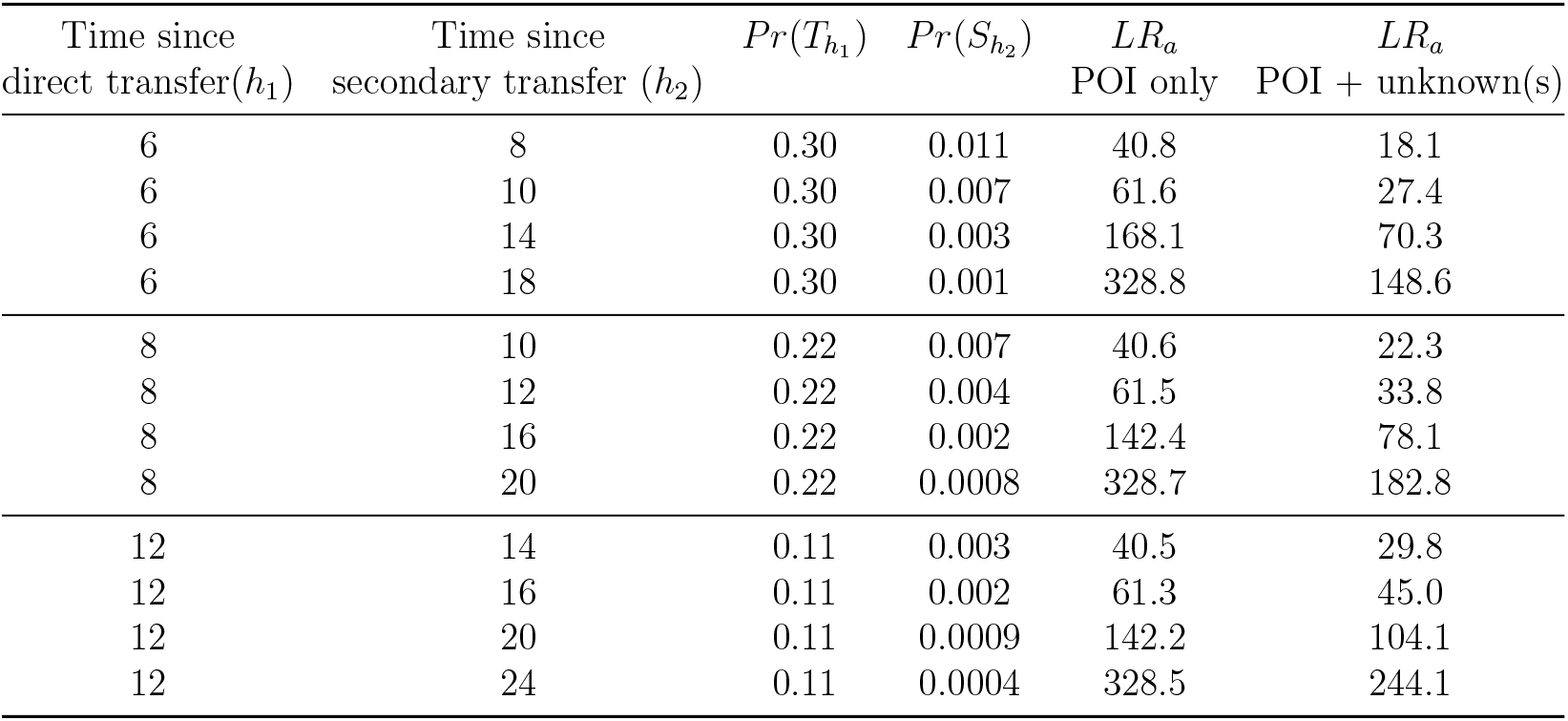
Activity level likelihood ratios from logistic regression where logistic regression decision threshold *x* = 3 and *n* = 1. All times (*h*) are measured from the time the sample was collected e.g. in the first row, the assault occurred six hours before sampling, and social contact at eight hours beforehand.

## 13. Multiple contacts and cohabitation

The previous experiment simulated the effect of a single contact. However this does not simulate the effects of transfer between individuals who spend a period of time in close contact with each other. This includes cohabitees and individuals who meet together in social settings where there is an opportunity for multiple secondary and/or direct transfer events.

As part of the case-circumstances, there needs to be a consideration of the potential for recovering DNA from (innocent) direct transfer, as well as for secondary transfer events.

### 13.1. Simulation of secondary transfer accumulation

Prevalent DNA from a partner of a cohabitee will build over a period of time. The amount that accumulates onto a skin surface will depend upon the number of contacts that individual *X* has with a surface bearing DNA of individual *Y*. The accumulation over a 24hr period is shown in the example (Fig. 5) where *Pr*(*S* |*h* = 0, .., 24, *n* = 1, .., 4, *x* = 3). A steady state is achieved where the DNA load that is gained balances the amount which is lost. This steady state occurs after a period of time (c. 12 hours).

**Figure 5.**
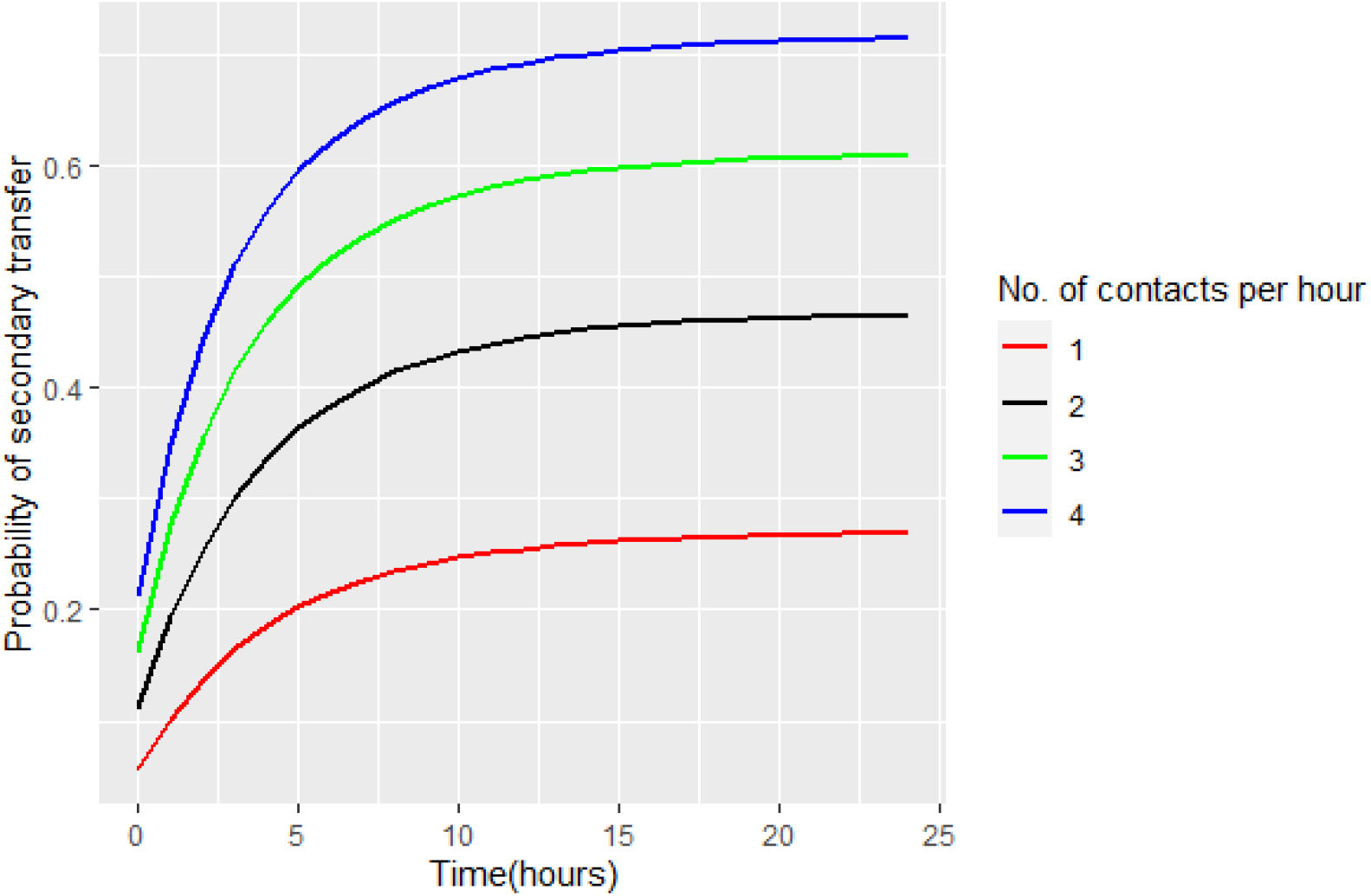
Probability of DNA recovery, given multiple secondary transfer events relative to time duration of exposure, where *log*_10_*LR*_*ϕ*_ > 3. A range of 1,..,4 contacts per hour are simulated up to 24 hours.

### 13.2. Sensitivity analysis (example)

As discussed earlier, the number of contacts per hour will be informed by case-circumstances. The question of their cumulative effect upon the likelihood ratio can be assessed by sensitivity analysis [1, 13] -where the values of parameters are altered to determine the effect upon the *LR*_*a*_. If the relative change in the magnitude of *LR*_*a*_ is small, it is said to be insensitive to the parameter values tested.

The alternative propositions are:

1. Victim was assaulted and had social interaction with X
2. Victim only had social interaction with X

#### Background information

The victim visited the suspect’ s home for a period of 12 hours. During this time there was no direct contact, although the victim made full use of the accommodation’ s facilities and had a meal/ drinks etc. According to the prosecution an assault occurred at time *y* and she left the premises and samples were taken from her skin by police at time *z*.

Table 5 illustrates activity level likelihood ratios achieved for a range of logistic regression conditioned sub-source likelihood ratios. The POI is accused of assault at time *y*. However, the individuals were in close contact with each-other (no direct contact in this example) for the period of time indicated. Note that this may be a cohabiting couple and the time interval where secondary transfer can occur can be considered to be the time since bath/shower of the victim, which can be assumed to remove all foreign DNA from the skin [14]. All times in Table 5 are referenced to the difference between time of sampling and collection *h*, except for the first example where the time of assault is given at time zero, *h* = 0, with cohabitation time interval as 0-12hrs. If the result of the assault results in the instant death of the individual, then there is no opportunity for further gain or loss of DNA -the clock stays at time zero, provided the body is undisturbed before discovery. The data show that for this set of circumstances, the *LR*_*a*_ is increased with ‘ POI only’ results, compared to ‘ POI + Unknown’. With the latter, the evidence weakly favours the defence proposition for low subsource *LR*_*ϕ*_ s, and weakly favours the prosecution proposition with increasing sub-source *LR*_*ϕ*_ s.

**Table 5:** Modelling multiple secondary transfer events which may occur between cohabitees. Likelihood ratios were recorded where *H*_*p*_ is conditioned on secondary transfer with *n* contacts per hour for a period of 12 hours and one direct contact at the time indicated in column 1. Column 2 is the time interval for *H*_*d*_: secondary transfer with *n* contacts per hour and the direct contact event by an unknown individual at alleged time of the assault. Four separate time intervals were tested as indicated. *H*_*p*_ and *H*_*d*_ propositions tested were ‘ POI only’ and ‘ POI and unknown’. The *LR*_*a*_ is always higher for the former

|  |  |  | POI only |  |  |  | POI and U |  |  |  |
| --- | --- | --- | --- | --- | --- | --- | --- | --- | --- | --- |
| Time of<br>assault<br>(since item<br>sampled) | Co-hab.<br>time interval | No of contacts<br>per hour | 1 | 2 | 3 | 4 | 1 | 2 | 3 | 4 |
| | | sub-source<br>$\log_{10} LR_\phi > x$ | | | | | | | | |
| 0 | 0,12 | 1 | 3.5 | 2.9 | 2.8 | 2.8 | 0.6 | 0.5 | 0.5 | 0.5 |
|  |  | 2 | 4.9 | 3.5 | 3.1 | 2.9 | 0.9 | 0.6 | 0.5 | 0.5 |
|  |  | 3 | 7.2 | 4.6 | 3.70 | 3.3 | 1.3 | 0.85 | 0.7 | 0.6 |
|  |  | 4 | 12.0 | 7.0 | 5.3 | 4.5 | 2.13 | 1.2 | 1.0 | 0.8 |
|  |  | 6 | 15.4 | 8.5 | 6.17 | 5.0 | 3.8 | 2.1 | 1.5 | 1.2 |
|  |  | 8 | 25.1 | 13.3 | 9.4 | 7.4 | 6.9 | 3.7 | 2.58 | 2.0 |
| 3 | 6,2 | 1 | 3.8 | 2.8 | 2.5 | 2.3 | 0.9 | 0.7 | 0.6 | 0.6 |
|  |  | 2 | 6.6 | 4.1 | 3.23 | 2.82 | 1.8 | 1.1 | 0.88 | 0.8 |
|  |  | 3 | 11.6 | 6.5 | 4.8 | 4.0 | 3.5 | 2.0 | 1.46 | 1.2 |
|  |  | 4 | 22.5 | 12.0 | 8.5 | 6.7 | 6.9 | 3.7 | 2.59 | 2.0 |
|  |  | 6 | 27.1 | 14.2 | 9.9 | 7.8 | 10.4 | 5.43 | 3.8 | 3.0 |
|  |  | 8 | 44.2 | 22.7 | 15.57 | 12.0 | 17.6 | 9.1 | 6.2 | 4.8 |
| 6 | 12,2 | 1 | 4.8 | 3.1 | 2.5 | 2.3 | 1.6 | 1.0 | 0.8 | 0.8 |
|  |  | 2 | 10.1 | 5.7 | 4.23 | 3.5 | 4.0 | 2.2 | 1.7 | 1.4 |
|  |  | 3 | 20.3 | 10.8 | 7.6 | 6.0 | 9.1 | 4.8 | 3.4 | 2.7 |
|  |  | 4 | 44.9 | 23.1 | 15.8 | 12.1 | 21.3 | 10.9 | 7.5 | 5.76 |
|  |  | 6 | 49.9 | 25.5 | 17.4 | 13.3 | 26.4 | 13.5 | 9.2 | 7.1 |
|  |  | 8 | 78.0 | 39.6 | 26.8 | 20.4 | 42.3 | 21.4 | 14.5 | 11.0 |
| 6 | 24,36 | 1 | 21.8 | 11.6 | 8.2 | 6.5 | 7.2 | 3.8 | 2.7 | 2.1 |
|  |  | 2 | 80.7 | 41.0 | 27.7 | 21.1 | 31.5 | 16.0 | 10.8 | 8.3 |
|  |  | 3 | 246.0 | 123.6 | 82.8 | 62.4 | 111.8 | 56.2 | 37.6 | 28.4 |
|  |  | 4 | 806.6 | 403.9 | 269.7 | 202.6 | 387.7 | 194.1 | 129.6 | 97.4 |
|  |  | 6 | 592.6 | 296.9 | 198.3 | 149.0 | 312.0 | 156.32 | 104.4 | 78.5 |
|  |  | 8 | 798.9 | 400.0 | 267.1 | 200.6 | 431.6 | 216.1 | 144.3 | 108.4 |

If *H*_*p*_ is true, then the *LR* increases as the time of ‘ innocent’ contacts becomes more distant from the assault. In casework, the frequency, type and duration of contacts will be assigned as a prior from careful consideration of the case-circumstances.

## 14. Discussion

### 14.1. The requirement for a framework to model likelihood ratios

Although there are many papers that describe various aspects of reporting activity level propositions (see reviews by Taylor et al [15] and Oorschot et al [16]), it can be difficult to extract the information needed to inform BNs, In this paper we have provided a framework to calculate activity level likelihood ratios based on experimental datasets of direct and secondary transfer.

Case-work circumstances are often complex; it may be necessary to model multiple contacts (Table 4). Oorschot et al [16], point out that little is known about accumulation of DNA on an object due to increased duration and frequency of contact. Goray and Oorschot [17] carried out an interesting study that highlighted the complexities of social interactions, clearly demonstrating that transfer events cannot be considered as ‘ one-off’. As an example they examined the social exchanges of three individuals having a drink together while sitting at a table; there were multiple opportunities for transfer within a 20 minute window where DNA was secondarily and directly transferred between objects and onto hands. They found: “In 29.2% of the samples, mixtures from two or three contributors (including the owner of the hand) and one other participant were obtained.” The authors don’ t provide likelihood ratios, but this observation is consistent with the simulations in our study Fig. 5), conditioning on *n* = 1 contact per hour over a 14hr period, where *Pr*(*S* |*h* = 0, .., 14, *n* = 1, *x* = 3) = 0.26. Oorschot et al [18] also carried out a separate study showing that individuals are very tactile, and will often touch objects and themselves over short periods of time

Our framework accommodates the requirement to model multiple contacts, including cohabitation, and it can be easily extended. We use the sub-source likelihood ratio (calculated by *EuroForMix*) as a ‘ quality indicator’, which simultaneously takes account of mixture proportion, average peak height expectation, and peak height variability. Sub-source *LR*_*ϕ*_ s were used to inform a modified BN described in supplement 1 of Gill et al [1]. This BN also takes account of background DNA from unknown individuals (Table S1) under the ‘ POI and Unknown’ outcome. It was shown that *LR*_*a*_s are much reduced when background is present compared to the ‘ POI only’ outcome.

The only comparable literature, relating to skin surface DNA transfer, persistence and recovery is by Bowman et al [2]. The authors carried out a study of direct transfer onto skin surfaces at *h*=0, 3, 24. Their demonstration utilised a count of self and non-self alleles. *LR*_*ϕ*_ s were also measured and it was shown that there were more instances of *LR*_*ϕ*_ > 1 when the pressure of contact was greatest. However, the results were quite varied: the authors showed *LR*_*ϕ*_ > 1 with “0.56 and 0.77 of samples at 0hrs with medium pressure and heavy pressure and friction respectively”. This corresponds to 0.51; 0.64; 0.76 (5 percentile; median; 95 percentile) in the study reported here i.e. our results are broadly consistent. The authors concluded that it was possible to obtain offender DNA 24hrs post assault, with one example of extremely strong evidence at this time interval. In our study, the highest *LR*_*ϕ*_ = 1.6 at 24hrs and 16.2 at 18hrs. But, as noted by us and Bowman, there is a limitation of small sample sizes.

Bowman also noted the importance of taking account of background DNA. This is also accommodated in our framework and a sensitivity analysis is described in section 14.7.

At 3hrs, from posterior logistic regression parameter, the proportion of observations *LR* > 1 was 0.53, which was similar to that recorded by [2] for their experimental scenario (B) at 0.55, but their average across all scenarios were much lower at 0.22. There were differences in experimental design and multiplex, for example, participants were asked not to wash the area of contact, whereas this was not controlled by Bowman. [2].

### 14.2. Methods of comparison

Various methods of comparison have been used in the literature: including counting self vs non-self alleles, mixture proportions and DNA quantities recovered [2, 9,19,20,21,22,23]. However, mixtures commonly include background DNA from unknown contributors which interfere with positive identification of POI alleles. Furthermore, high counts of POI alleles do not necessarily coincide with *LR* > 1.

Using *EuroForMix*, conditioning upon ‘ *H*_*p*_ is true’, it is possible to extract mixture proportion, *M*_*x*_, average peak height (*Pk*_*ave*_) and peak height variance (*Pk*_*var*_). A POI centred (pseudo DNA quantity) measure can be created by multiplying together *M*_*x*_ × *Pk*_*ave*_. However, in our hands, this method did not seem to perform as well as the sub-source likelihood ratio (*LR*_*ϕ*_) (data not shown).

The following critique was kindly provided by Duncan Taylor (pers. comm.):

*“I was surprised to hear that the use of DNA amounts performed worse than using LRs for modelling when transfer is considered to have occurred. I agree that measures such as numbers of alleles or mixture proportions will not perform well as they are summaries of the data that lose information, but the LR should really be a reflection of the amount of DNA, but with additional confounding effects that are difficult to account for. For example your model will be affected by the rarity of someone’ s profile i*.*e. a rare profile will lead to a higher transfer probability, but of course this is just an artefact of the model, and not a real-world effect. The other aspect of using the LR is that the presence of other DNA can effect transfer probability i*.*e. a person may have a probability of transferring some DNA to an item from touching, say 1ng. You sample the item, obtain a single source profile with a high LR and this is counted as a transfer. But now imagine that the person has other DNA on their hands, they still transfer 1ng of their own DNA, but also 1ng of another person’ s DNA. Now faced with an even 2 person mixture the LR will be greatly reduced. It could result in the event no longer being counted as a transfer (if the LR drops below a threshold), even though the person has transferred the same amount of DNA both times. I have always thought that using DNA amounts is preferable for these reasons and that because DNA extraction sits prior to PCR, electrophoresis and interpretation it means that laboratories can use the same model, even if these downstream laboratory processes differ from the published study*.*”*

Therefore, we accept that ideally a method is needed that incorporates (a selection of): peak heights, the number of alleles, quantity of DNA, mixture proportions, but not allele frequencies. The method should be conditioned upon the POI for the numerator, but not the denominator. Currently, we are not sure how best to achieve this. and is consequently the next challenge to pursue. Fortunately, any set of data, can be plugged into the existing ALTRaP program, no matter how derived, hence this will facilitate future studies.

### 14.3. Data limitations: problem of rare observations

We have provided a framework to interpret recovery of DNA from skin to skin contact where secondary/direct transfer is conditioned. This model is based upon a logistic regression of direct transfer data and relies upon the reasonable assumption that secondary transfer persistence and recovery follows the same decay curve. It is much easier to provide data for direct transfer studies because positive results are common within the first 12 hours of deposition. To construct such logistic regression curves, it is important to design experiments that cover a sufficient range of time so that a spread of data are achieved. At 24hrs, observation of positive data is problematic because probabilities of DNA recovery are low and sample sizes are relatively small -this is ameliorated by the logistic regression line which is in part predicated by data observations *<* 12hrs. The projection of the line can continue beyond 24hrs, but the variance increases.

Early methods e.g. Ladd et al [24] failed to detect any secondary transfer, multiplexes were not as efficient as those currently utilised. Even with modern multiplexes, the event of secondary transfer is poorly characterised. Consequently, recovery of DNA following secondary transfer is comparatively uncommon -probabilities are typically *Pr <* 0.01, which means that large sample sizes are required in order to assign probabilities. To maximise the number of observations, we carried out data analysis where the time difference between deposition and sampling was 1hr or less. The point corresponding to the position on the direct transfer logistic regression curve was assigned by calculation of an offset value (*f*_*x*_) and adjusted so that *h* = 0 (Fig. 3).

### 14.4. Modelling multiple contacts with ALTRaP

The framework provided here allows for multiple direct and secondary transfer events and includes simulation of cohabitation (Fig. 4). In casework, it may be difficult to inform the number and kinds of contacts, hence sensitivity analysis of a range of reasonable options may be called for [1, 13]. In order to carry out the analysis, the program ALTRaP (Activity Level Transfer, Recovery and Persistence) was written in R-code which is open-source and freely available at: https://activitylevel.shinyapps.io/shinyaltrap/. The source code and user manual can be accessed at: https://sites.google.com/view/altrap/home, along with the raw data of direct and secondary transfer used to create the tables and figures in this paper. ALTraP provides a basis to compare independently prepared data in order to facilitate comparative studies, as well as promoting a standardised analytical framework.

### 14.5. Reporting activity level propositions

As Champod [25] points out: the landscape is complicated and there are two important questions:

1. Is it the role of the scientist to offer guidance as to the probability of the DNA findings given various transfer mechanisms put forward by the parties depending on the case circumstances?
2. Can a forensic scientist robustly assess the probability of the DNA findings given alleged transfer scenarios in the current state of knowledge?

There is a strong support for view (1) in the ISFG DNA commission documents [1, 26]. In order to provide a ‘ robust’ assessment of the evidence, this requires the formulation of probabilities that must be input into a statistical framework that takes full account of the case circumstances from both the prosecution and defence perspective.

There will always be uncertainty relating to case-circumstances. This is why it is necessary to carry out exploratory analysis of reasonable assumptions that are put forward by the defence and prosecution cases. In the absence of complete information, the scientist can carry out exploratory and sensitivity analysis to discover the effect on the likelihood ratio of varying those parameters believed to be relevant to the case.

Once candidate model(s) are decided, then the issue remains about how to report the findings in accordance with Champod’ s second requirement which asks whether findings can be *robustly* reported? In their ‘ position paper’, Taylor et al [13] define robustness as: “our evaluation will be considered robust, if the system, informed by a different set of experiments, leads to similar *LR* values.” and “Sensitivity analysis is achieved by simulating cases under a range of datasets and exploring the impact of this on the output.” This theme was further explored by Samie et al. [27] who used simulation to determine those parameters that are most sensitive to data paucity, and require focused research effort.

There are two different aspects to sensitivity tests: first is the effect on the likelihood ratio when different parameters are chosen e.g. number of *n* contacts per hour. The second aspect addresses the variability of the *LR*_*a*_ once *n, T*_*h*_, *S*_*h*_ are fixed.

Here we used Bayesian logistic regression analysis to analyse the data. Each test results in 4000 pairs of data-dependant simulated coefficients that are used by the logistic function in order to generate likelihood ratios. By way of example, Fig. 6 is generated from the *T*_*h*=6_, *S*_*h*=18_, *n* = 1 plot shown in Table 4. For each value of logistic regression decision threshold *x*, a density (violin) plot is shown. Superimposed in green, is a box-whiskers plot, and behind, the blue rectangle delimits 0.05 and 0.95 percentiles, whereas the red rectangle delimits 0.025 and 0.975 percentiles. This plot was generated by the ALTRaP (tables can also be generated).

**Figure 6.**
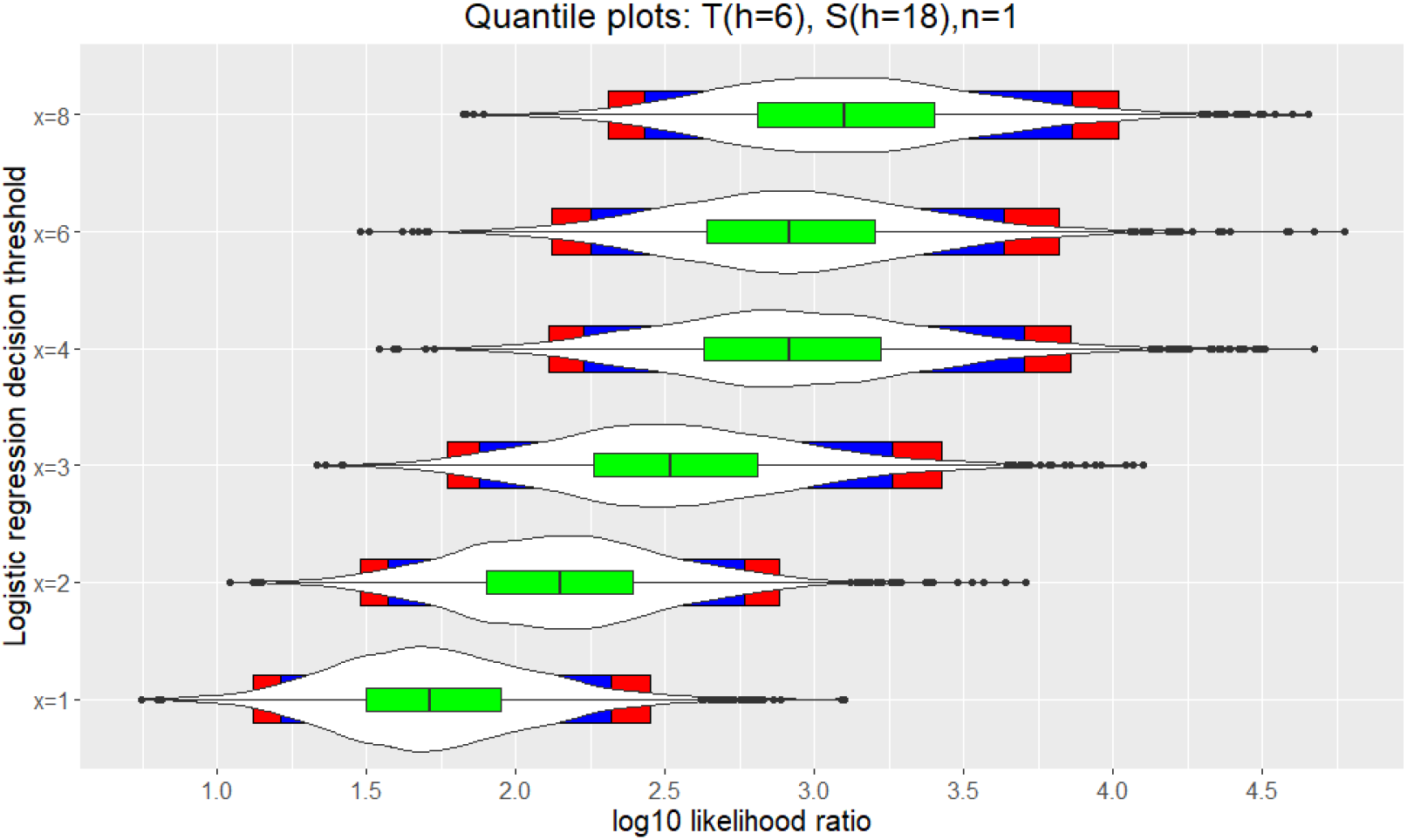
A total of 4000 *log*_10_*LR*_*a*_s, per value of *x*, simulated from logistic regression coefficients using *T*_*h*=6_, *S*_*h*=18_, *n* = 1. For each value of logistic regression decision thresh-old *x*, a density (violin) plot is shown in white. Superimposed is a box-whiskers plot in green, and behind, the blue rectangle delimits 0.05 and 0.95 percentiles, whereas the red rectangle delimits 0.025 and 0.975 percentiles.

### 14.6. Calculating a plausible range of probabilities of secondary transfer from small datasets

In section 6.3 the issue of small datasets was highlighted in association with the assignment of probabilities of secondary transfer. Data collected from zero -1 hour after the transfer experiment, were subject to bootstrapping [28] in order to generate 1000 new randomised sets of samples. For each dataset, new Pareto distribution parameters were calculated, and *Pr*(*S* |*x*) was determined as describe in section 8. Quantiles were calculated ranging between 0.5-0.99 (Table 6). Finally, for given values of *x, n*, ALTRaP can be used to display the variance of likelihood ratios based upon a plausible range of *Pr*(*S*) values from the bootstrap experiments. Fig. 7 uses the same parameters as those used to generate Fig. 6, and illustrates quantile plots for *x* = 3, *n* = 1. With this example, the plausible range of *Pr*(*S*) = 0.05 − 0.12 is defined by 0.5-0.99 quantiles. This has only a small effect on the median *log*_10_*LR*_*a*_ which drops from 2.5 -2.1. In conclusion, the impact of small data-sets and low positive observations, can be assessed by sensitivity analysis and is therefore not a limitation.

**Table 6:** Summary statistics from 1000 bootstraps of the secondary transfer data (*h* = 0 and 1 hours combined), The plausible range is represented by the 0.5 -0.99 quantiles. *x* is the logistic regression decision threshold

| $x$ | Quantile | | | |
| --- | --- | --- | --- | --- |
|  | 0.50 | 0.75 | 0.975 | 0.99 |
| 1 | 0.164 | 0.203 | 0.269 | 0.292 |
| 2 | 0.089 | 0.112 | 0.161 | 0.186 |
| 3 | 0.052 | 0.065 | 0.105 | 0.124 |
| 4 | 0.032 | 0.040 | 0.070 | 0.089 |
| 6 | 0.014 | 0.017 | 0.034 | 0.046 |
| 8 | 0.007 | 0.009 | 0.023 | 0.028 |

**Figure 7.**
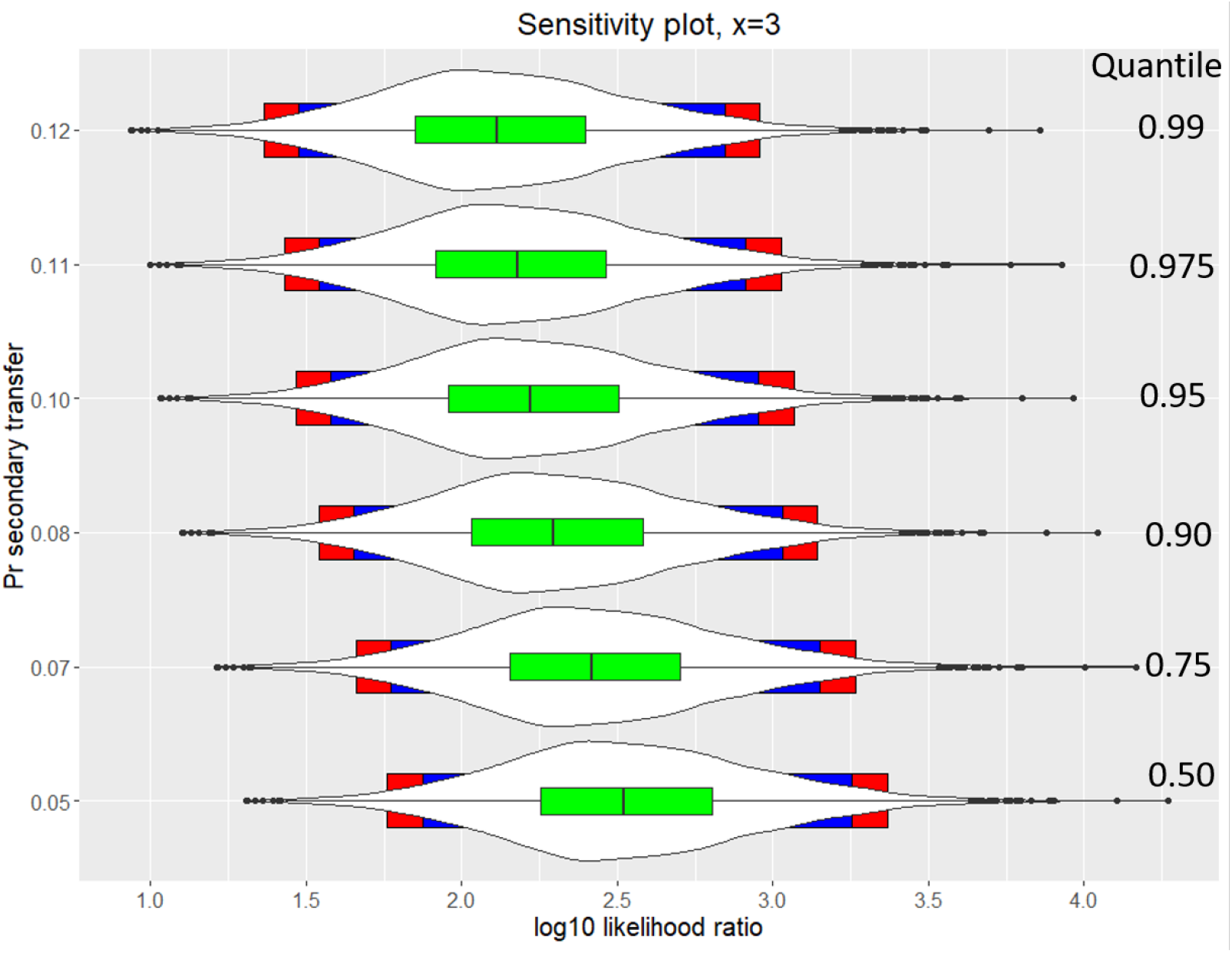
A total of 4000 *log*_10_*LR*_*a*_s simulated from logistic regression coefficients using *T*_*h*=6_, *S*_*h*=18_, *n* = 1, *x* = 3. The quantiles of *Pr*(*S*) shown on the *y* −axis are based upon analysis of 1000 bootstraps of the secondary transfer data (*h* = 0, 1). For each value of logistic regression decision threshold (*x*), a density (violin) plot is shown in white. Superimposed is a box-whiskers plot in green, and behind, the blue rectangle delimits 0.05 and 0.95 quantiles, whereas the red rectangle delimits 0.025 and 0.975 quantiles.

### 14.7. Sensitivity analysis of background DNA

Background is defined in section 3.2 as: “DNA present from unknown sources and unknown activities”. The likelihood ratio eq. 2 is calculated with one or more unknown contributors in the numerator and denominator. In our study we used a level of *Pr*(*B*) = 0.36 throughout. This was an average taken across direct and secondary transfer experiments. If the data were analysed separately, the probabilities were *Pr*(*B*)= 0.25 and 0.59 respectively. To examine the sensitivity of the *LR* to the background parameter, a range of probabilities were input into ALTRaP and the results are shown in Table 7.

**Table 7:** Comparison of *log*_10_*LR*_*a*_s where *S*_*h*=18_, *T*_*h*=6_, varying probability of background *Pr*(*B*). There is no effect when the ‘ POI only’ is recovered, along with prevalent DNA from ‘ known’ individuals. The LR is always lower when background is present. The *LR*_*a*_ increases as *Pr*(*B*) increases.

| Direct | Secondary transfer |  |  |  |  |
| --- | --- | --- | --- | --- | --- |
|  | Probability of background |  |  |  |  |
|  | 0.1 | 0.2 | 0.36 | 0.8 | 0.99 |
| 1.71 | 0.8 | 1.05 | 1.23 | 1.43 | 1.47 |
| 2.15 | 1.33 | 1.57 | 1.74 | 1.92 | 1.96 |
| 2.52 | 1.79 | 2.02 | 2.17 | 2.33 | 2.36 |
| 2.92 | 2.22 | 2.44 | 2.59 | 2.74 | 2.77 |
| 2.92 | 2.3 | 2.51 | 2.64 | 2.77 | 2.8 |
| 3.1 | 2.5 | 2.7 | 2.84 | 2.96 | 2.98 |

Background has no effect on the *LR*_*a*_ for the ‘ POI only’ result. The ‘ POI and unknown(s)’ *LR*_*a*_ is always lower, but as the *Pr*(*B*) increases, so does the *LR*_*a*_. The biggest effect on the *LR*_*a*_ is the detection of background DNA itself. The overall effect of the *level* of background on the *LR*_*a*_ is small. Background levels on skin recorded by other authors are quite high e.g. [2], their Fig. 1, where *>* 10 non-self alleles are found in *>* 50% of their experiments. Graham et al. [29] recorded 23% from necks of simulated strangulated victims and 31.6% from children [30] (note that these studies were carried out several years ago with less sensitive multiplexes). These findings support a proposition that background levels of DNA are relatively high on skin.

### 14.8. Reporting the evidence at activity level

In the worked examples (sections 11 and 12), we stressed the importance that the evidence at sub-source level was accepted by the court, otherwise it cannot be evaluated at the activity level. It is the responsibility of the scientist to make this clear in his/her report by including a statement along the lines outlined by the ISFG DNA Commission [26]:

*“Assuming it is accepted by all parties that the origin of the DNA is Mr X, the probability of recovering DNA on this item, with the observed relative quantity/quality, given that the person performed the alleged activity is*… *“*.

The scientist provides the information and the caveats. There is reliance placed upon the lawyers and the judiciary to ensure that the necessary steps are followed to establish that agreement does indeed exist between the relevant parties on “the origin of the DNA”. An informative example where the procedure was properly followed is provided by an appeal court of England and Wales, Regina v. Weller [31], where in the judgement it was stated:

*“It therefore was common ground at the trial and on this appeal that the DNA had come from Emma*.*”*

As the source of the DNA was not contested, the court moved on to discuss the next level in the hierarchy of propositions, i.e. the activity level. To reiterate Champod [25] “the landscape is complicated”; it is pertinent to ask how the court may be helped without becoming impeded with details that may be difficult for the lay-person to understand? In England and Wales, a judge may direct scientists acting on behalf of the defence and prosecution to prepare a joint statement that clearly states the areas of agreement and disagreement [32]. The advantage is that the debate between scientists becomes *collaborative* rather than *combative*. Once the common ground is made clear, the court is able to focus energy on any disagreements that may exist. In support, a view eloquently put by the appeal court of England and Wales, Regina v. Weller is as follows:

*“It is, we think, becoming increasingly common for there to be little dispute in the majority of cases as to whether the DNA is the DNA of a particular person, although that may be in issue where quantities are very small or the amount obtained has been difficult in analysis or there are mixed profiles. But where, as in this case, that is clear, it is essential that this issue is put before the jury as admitted and agreed expert evidence. It makes the task of the jury so much easier if they do not have to plough through and listen to evidence that is simply not in dispute. It enables the jury to perform its essential function of assessing, where it is agreed that there is a sufficient scientific basis for expert evidence to be given, what is in fact in issue without being troubled by matters that are not*.*”*

Indeed, the dangers of *not* following such an approach are manifestly illustrated by a remarkable case that illustrates the dangers of inadequate representation at activity level. From the appeal court of England and Wales, Queen v. William Francis Jones [33]: a DNA profile was found on the pin of a hand-grenade and a statement was provided at the original trial:

*“the conclusion was that the mixed DNA result was 1 billion times more likely if the DNA came from the appellant and two unknown and unrelated persons”*.

This evidence was sufficient for conviction. However, at appeal, the experts were directed by the judge to prepare a joint statement. There was agreement about the value of the evidence at sub-source level, but at the original trial, the activity level had not been properly addressed. It was agreed that this had been a serious omission. If a jury is told that the sub-source evidence is an impressive *LR*=1bn, in the absence of subsequent discussion about the limitations of the evidence, then it is highly likely to result in a wrongful conviction.

Because of lack of data or a model, the experts were unable to help the court assess the probabilistic value of the evidence at activity level. Consequently, the conviction was overturned. The court-ruling is available at [33] and details are paraphrased in supplement S6.

In a wider context, this case shows is that our current state of knowledge to provide probabilistic solutions to the questions posed by typical casecircumstances is limited. Once models are developed and the necessary data generated, then they become ideal platforms to facilitate constructive debate between the defence and prosecution scientists within a collaborative framework. Different kinds of transfer, persistence and recovery events may be modelled. hence the procedure becomes exploratory. Not only do defence and prosecution scientists need training to carry out evaluation at activity level, thereby creating a level playing field, but the court environment also needs to be fit for purpose to allow constructive debate to occur [34].

### 14.9. Reporting the likelihood ratio

The question remains how to report the evidence? Either a range of *LR*_*a*_s could be reported or some lower limit corresponding to to a percentile (e.g. 0.025). Alternatively a median could be used. For example, for *x* = 3 in Fig. 6, the range of *LR*_*a*_ defined by 0.025 and 0.975 quantiles is approximately 60-3000 with the median at 400. The range of *LR*_*a*_s values gives assurance that the assessment is robust because it is within ‘ acceptable’ limits (the scientist decides what is ‘ acceptable’: for a comparison, see Fig. 10 of Taylor et al [13]). The reader will note that in this paper we have used single *LR*_*a*_ values (medians based on the posterior distribution of logistic regression coefficients) throughout, per experiment. This practice accords with the recommendations of Taylor et al. [13] who state: “This single value (generally an order of magnitude will suffice) takes into account the data and the knowledge that we have: it encapsulates all our uncertainty.” Also see Taroni et al. [35] for further compelling arguments related to reporting a single value.

During a review of an earlier transcript of this paper, a referee wrote:

*“I have always subscribed to the Taroni view -a single order of magnitude is what should be reported. But perhaps we could do more to assist the scientist here. For example, in case A the median is 400 and the 95 interquantile range is 60-3000; in case B the median is 400 and the interquantile range is 500-700. Does the scientist report exactly the same in both cases? Some assistance in the form of a facility to integrate over a meaningful distribution would seem desirable*.*”*

Probabilities are personal (ENFSI Guideline,note 3 [12]) and Biedermann et al. [36].The ENFSI guideline also emphasises the need for transparency -the practitioner must disclose the source of the data and software used to make inferences so that it is open to challenge. The advantage of ALTRaP is that it provides a framework to promote discussion -either new data or new parameters can be explored to determine the effect upon the *LR*_*a*_. This is particularly useful when the experimental data-size is limited. The plots illustrated in Fig. 6 should also be made available for disclosure. It is neither realistic nor necessary for models to be ‘ perfect’. Over time, as new information is gained, models will evolve to take account. For example, shedder status will doubtless have an effect on the results, but here it is not currently modelled. All models are based upon *existing* knowledge. Improvements take account of more information, and the new model is preferred as a result, but this does not negate earlier results based upon fewer assumptions.

Finally, we note that there is a paucity of data in the literature. Studies would be greatly facilitated if datasets were readily available to the community, along with standardised methods of experimental design and analysis [34]. This is important because the issue of reproducibility of data-collections needs to be addressed. Similar pleas have also been made by Meakin and Jamieson [37]. Gosch and Courts argue that: “available literature lacks quality and systemization”.

### 14.10. Inter-laboratory comparisons

It would be possible to make comparisons with different multiplexes using different populations. To carry out inter-laboratory comparisons, data can be rescaled in order to be directly comparable with the results of this study. To do this, the laboratory should calculate the average sub-source likelihood ratio for full profiles of the multiplex, population with *Fst*. Calculate the rescaling factor *q/z* where *q* is the average *LR*_*ϕ*_ of Fusion 6C multiplex and Norwegian frequency database, and *z* is the average *LR*_*ϕ*_ of the new multiplex/population database/*Fst* under investigation. Each individual (*i*) sub-source *LR*_*ϕ*_ must be multiplied by the rescaling factor before carrying out logistic regression analysis. Using *log*s: *log*_10_*LR*_*ϕ q*_ = 32.8; hence *log*_10_*LR*_*ϕ rescaled*_ = *log*_10_*LR*_*ϕ i*_ + 32.8 *− log*_10_*LR*_*ϕ z*_

### 14.11. An outline of future plans: introduction into casework

The intention is to develop new Bayesian Networks to address different activity level problems, ultimately providing a suite of programs. Before introducing into casework three steps are required:

1. More work is required to compare different methods of data presentation as outlined in section 14.2.
2. Collaboration between laboratories to generate sets of open-access data will enable comparative studies (we note that there is a lot of interest in the community for inter-laboratory exercises).
3. Users will need to be trained in both theory and practice.

ALTRaP is designed to be very easy to use, since no knowledge of computing is needed. It is very useful for training and demonstration purposes.

## 15. Conclusion

The most important effect on the likelihood ratio is time-dependancy of the probability of recovery of DNA following its transfer and persistence. When DNA is initially applied to a surface, it will dissipate or degrade, until a point is reached where it cannot be detected. The process is *continuous*. Over time, DNA is dissipated by activities such as rubbing against surfaces, or washing. We call this ‘ DNA decay’. Previous modelling of DNA decay has been discrete: e.g. Bowman et al [2] model transfer at 0, 3, 24hr. In contrast, our experimental design involved collection of data from multiple time points up to 24hr, so that a continuous logistic regression model could be applied to model the decay of DNA over time.

Secondary transfer is more difficult to model because it tends to be a rare event, particularly if the time period in question is lengthy. Our solution to this was to assign the probability of secondary transfer at time zero (using a Pareto distribution), followed by determination of the corresponding position on the direct DNA logistic regression. The advantage is that the curve of single logistic regression can describe both direct and secondary transfer for a given value of *x*.

We have used the sub-source likelihood ratio in all our modelling. In the context of activity level, previous authors have investigate DNA concentration and mixture proportion [38], and/or the number of alleles present in the sample attributed to the POI. Probabilistic genotyping also incorporates mixture proportion, DNA concentration (in the form of peak height) and alleles into a sophisticated model that generates the sub-source likelihood ratio (see deliberations at section 14.2).

Finally, if activity level propositions are to become widespread practice, user friendly models need to become available. Here we have provided such a model (ALTRaP); the program can be accessed from: https://activitylevel.shinyapps.io/shinyaltrap/ and opens in the browser window; the source-code for the R-environment, and user manual is available at https://sites.google.com/view/altrap/ and https://github.com/peterdgill/altrap. The Bayesian network is generalised, meaning that it can be used to describe many different kinds of events where secondary and direct transfer on skin surfaces are of interest; examples are provided in section 11.

### To summarise

1. The experimental design can be both simplified and standardised. We modelled direct transfer with a total of 88 samples. Determination of probability of secondary transfer at time zero was achieved with 23 samples.
2. The output of the sub-source *LR*_*ϕ*_was modelled to produce the activity level *LR*_*a*_
3. The model is simplified because it utilises a single logistic regression curve (dependant upon the value of *LR*_*ϕ*_) for all subsequent calculations.
4. The model forms the basis of ALTRaP, a user friendly program that has been generalised, and is intended to act as a bridge to encourage the evaluation of activity level evidence in court
5. Our experimental design is simplified, and precisely described so that it may be standardised. It is coupled with a method of analysis that can be easily adapted to incorporate different sets of data. Consequently, this approach will facilitate inter-laboratory comparisons. Those interested in collaboration should contact the authors as we are happy to make necessary changes to ALTRaP to improve its functionality.
6. To report, the median likelihood ratio should be used. Case-specific sensitivity analysis can be carried out with ALTRaP, to model parameter uncertainty, using quantiles.
7. The method is limited only to those cases where the prosecution proposition at sub-source level is understood to be true by the court. It cannot be used otherwise.

## Supporting information

supplement 8 secondary transfer data

Supplement 7 direct transfer data

## Acknowledgements

We are grateful to Duncan Taylor and an anonymous referee for their constructive comments. Also to Ingebjørg Heitmann for her technical assistance with the direct and secondary transfer experiments.

## Supplementary material

### S1. Derivation of formulae utilised in Bayesian Network

#### S1.1. Outline of the Bayesian Network

An outline of the generalised Bayesian Network is shown (Fig. S1, based upon Gill et al. [1], supplement 1, where a detailed explanation can be found for further details.

**Figure S1.**
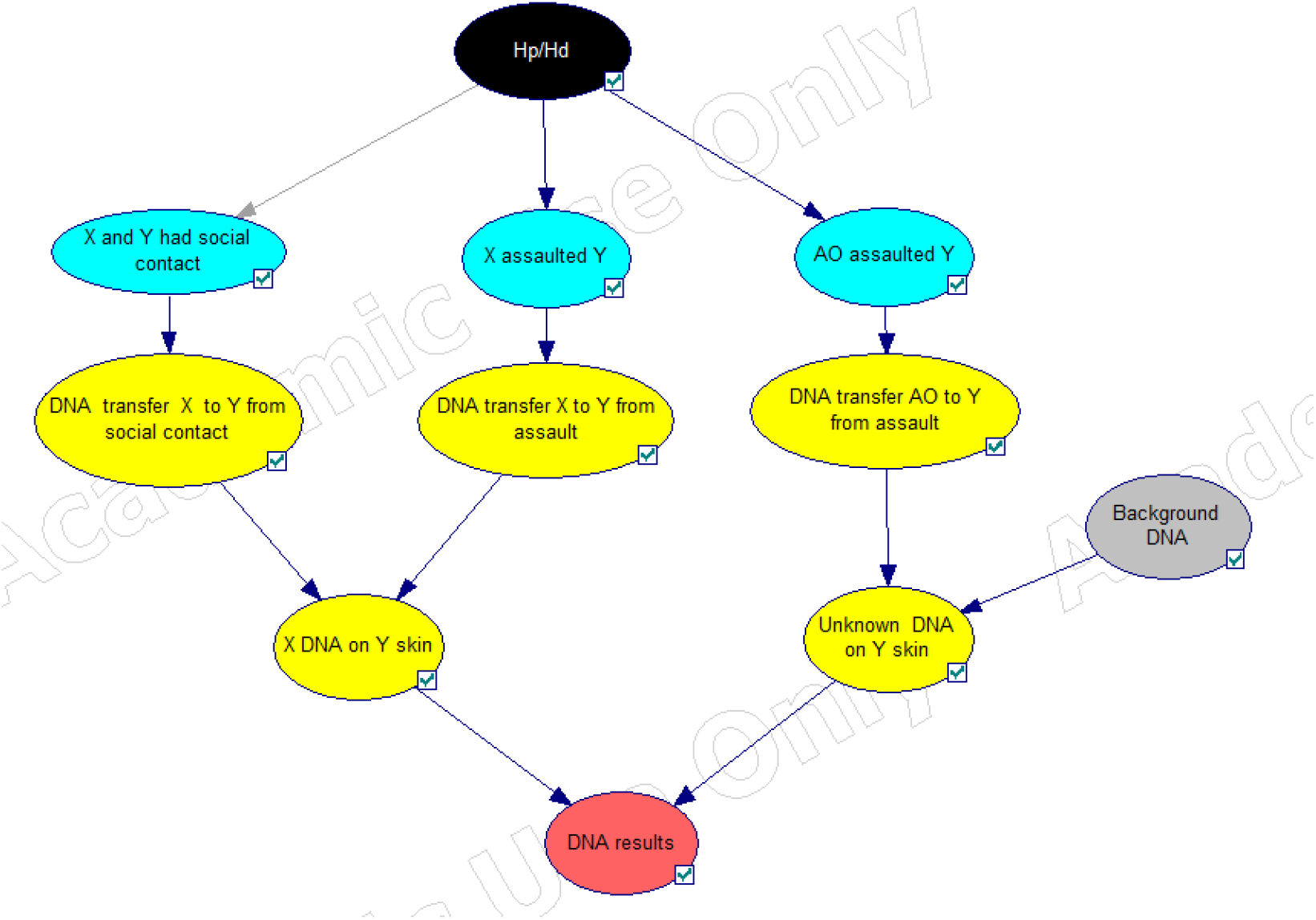
Generalised Bayesian Network used to evaluate the DNA results programmed into ALTRaP, given the activity level propositions summarising prosecution and defence’ s view of events. Nodes have been coloured so that black represents propositions, blue represents the sub-activities: social contact is modelled as common ground (*Pr* = 1 for both *Hp* and *H*_*d*_); “X assaulted Y” is proposed by *H*_*p*_ (*Pr* = 1); “AO assaulted Y” is proposed by *H*_*d*_ (*Pr* = 1). Yellow represents transfer and accumulation probabilities, grey represents a background node and red represents the results node. First layer of yellow nodes employ the word ‘ transfer’ in a broad sense – fully written: “… transfer from X, persistence and recovery from Y….”

#### S1.2. Probability of recovery of DNA from an unknown contributor (H_d_)

Conditioning that DNA is recovered from an unknown contributor(*U*_*d*_) is present under *H*_*d*_, this could come either from an unknown assailant or as a result of background (from one or more contributors) or both:

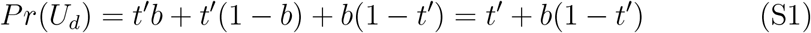

Where *t*^*′*^ is the probability of direct transfer, persistence and recovery from an unknown assailant and *b* is the probability of detecting background (always defined as DNA from *unknown* contributors); *Pr*(*U*_*d*_) is the combined probability of recovering unknown DNA. Under *H*_*p*_, *t* is the probability of transfer, persistence and recovery of DNA from the POI. There is no alternative assailant, hence the source of an unknown(s) can only come from background, hence *Pr*(*U*_*p*_) = *b* (*b* is common to *H*_*p*_ and *H*_*d*_). We introduce parameter *s* (common to *H*_*p*_ and *H*_*d*_) as the probability of secondary transfer, persistence and recovery.

Under *H*_*p*_, The probability of obtaining a DNA profile (*E*) only contributed by the POI, after direct transfer, with no background becomes:

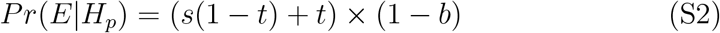

The probability of obtaining a DNA profile (*E*) contributed by the POI, with background from one or more unknowns:

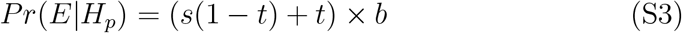

Under *H*_*d*_, direct transfer has not occurred from the POI, hence the probability of obtaining a DNA profile (*E*) only contributed by the POI, after secondary transfer, with no background becomes:

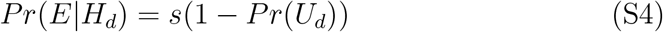

The probability of obtaining a DNA profile (*E*) contributed by the POI, with background from one or more unknowns:

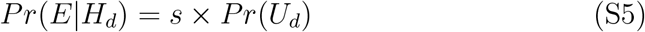

#### S1.3. ‘ POI only’ is observed

Note that prevalent DNA i.e from the ‘ victim’ is usually present but this does not affect the calculation.

Under *H*_*p*_, there was no alternative assailant and there is no background, hence *Pr*(no background) = 1 −*b*. Under *H*_*d*_, the POI is innocent as there is an alternative assailant -hence the absence of an unknown contributor is because of concurrent no background transfer and no assailant transfer *Pr* = 1 *− U*_*d*_

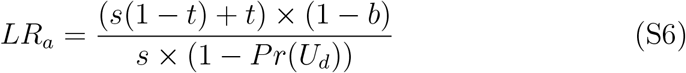

#### S1.4. ‘ POI and unknown’ is observed

Presence of prevalent DNA does not affect the calculation.

If unknown contributors are present then this can only occur from background under *H*_*p*_ with probability *b* and from either background and/or an unknown assailant under *H*_*d*_ with probability *s × Pr*(*U*_*H*_*d*)

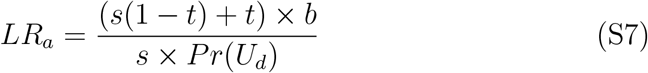

### S2. Derivation of formulae for computing purposes

A flexible program: Activity Level, Transfer, Recovery and Persistence (ALTRaP) was prepared using simple rule-sets. These are outlined in Table S1.

**Table S1:**
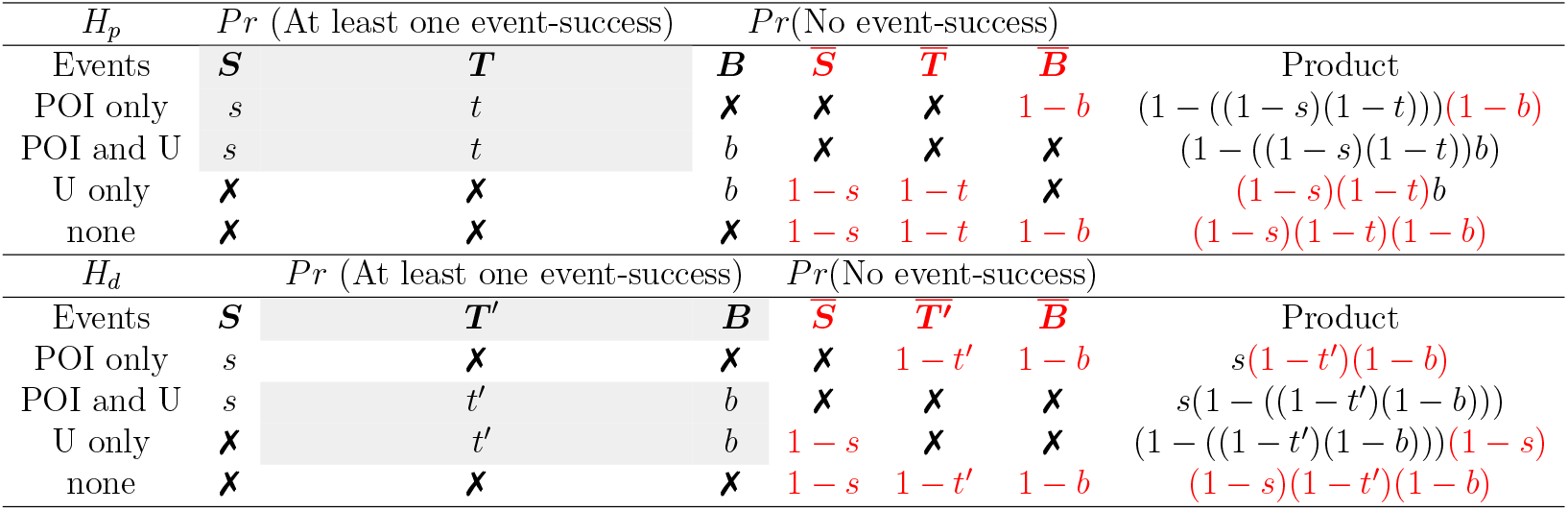
Derivation of formulae used to calculate likelihood ratios. Where the outcome is explained by event-successes eg. *S* or *T*, or no event-success is indicated by 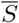 and 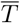 in red-type. There are three assignments per row along with three impossible assignments marked with X. Products are calculated using binomial expressions wherever the explanation of an observation could be from two or more events where ‘ one or more event-successes’ is required. Binomial expressions have grey background and the parameters differ between *H*_*p*_ and *H*_*d*_

The results listed in the first column are all possible outcomes, but their explanations are different under *H*_*p*_ and *H*_*d*_. Under *H*_*p*_, if DNA from the POI, is recovered, then it must have arisen either by direct transfer with probability (*t*) and/or by secondary transfer with probability (*s*), along with background (*b*) from one or more unknown contributors. Under *H*_*d*_, the POI is innocent and this can only be explained by secondary transfer (*s*); instead, an unknown perpetrator was responsible for the direct transfer event and his/her DNA was not recovered with probability (1 −*t*^*′*^). Background DNA will also result in the presence of DNA from unknown contributor(s). Under both *H*_*p*_ and *H*_*d*_, probability parameter *b* is in common. The complementary probabilities are:

1. No secondary transfer: 1 *− s*
2. No direct transfer from POI: 1 *− t*, or from unknown assailant: 1 *− t*^*′*^
3. No background DNA: 1 *− b*

The first step is to assign the events of *S, T, T* ^*′*^, *B* in terms of ‘ at least one observation’ or as ‘ no observations’ to each row of the table, If an event is impossible e.g. if DNA from ‘ POI only’ is recovered, then background DNA is absent and is assigned a probability of ‘ no observation’ 1 *— b*, and as the complement, the probability of ‘ one or more observations’ (1 − (1 −*b*) = *b*) cannot occur, it is marked X and takes no part in the calculation.

The second step is to combine the probabilities. Where events can happen either separately or together, they are binomially distributed, indicated by grey background in table S1. For example, in the row: *H*_*p*_ ‘ POI only’, either secondary and or direct transfer is responsible for the observation of the POI. The probability of event-success leading to ‘ one or more observations’ is 1*−*((1*−p*_*a*_)*×*(1*−p*_*b*_)…), whereas the probability of ‘ no observations’ is just (1 *−p*_*a*_)(1 *− p*_*b*_)… The products of each row are built from the probabilities of three elements: observations from a single event e.g. with probability *b*, one or more binomially distributed success-events from two or more parameters (1 *−* (1 *− b*)(1 *− t*)), or unobserved events (1 *− b*). Likelihood ratios are calculated from the resultant *H*_*p*_ and *H*_*d*_ probabilities.

#### S2.1. Extension

Identical results will be obtained with a formal Bayesian network (BN) such as Hugin, and this has the advantage of avoiding the need to derive formulae. However, with ALTRaP, 4000 different activity level likelihood ratios are simulated per experiment. Although this could be achieved computationally using an existing software like RHugin [39], chapter 3, it is much faster to use coded formulae. It is also an efficient way to expand the BN if more events are to be considered. For example, for the ‘ POI only’ category, we may propose two secondary transfer events at different times (*S*_1_, *S*_2_) and a single innocent direct transfer event (*T*_2_) in addition to the crime-related direct transfer (*T*_1_). The probabilities under *H*_*p*_ of one or more binomially distributed success-events and can be calculated as: 1 *−* ((1 *− s*_1_)(1 *− s*_2_)(1 *− t*_1_)(1 *− t*_2_)), where (1 −*s*_1_) is the probability of no event.

If the POI’ s DNA profile is observed, it is not possible to determine which event(s) were responsible as their respective results are indistinguishable. Consequently, with the above example, we estimate the probability of at least one event-success from the two secondary and two direct transfer opportunities. Because the calculation is repetitive, it is easy to program any number of direct and secondary transfers.

### S3. Pareto distribution fitting to secondary transfer data

The proportion of data (*k*) where *log*_10_*LR*_*ϕ*_ *>* 0.1, were excluded from the initial distribution fitting (fig S2) and proportion (*k*) = 0.67. This greatly improved the fit without affecting probabilistic calculations of *Pr*(*log*_10_*LR*_*ϕ*_ ≥*x*), since the logistic regression decision threshold *x* is always greater than or equal to one. The calculation has to be rescaled with respect to the proportion *k*:

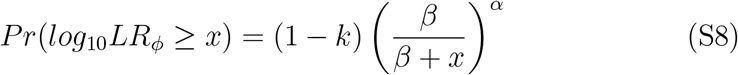

Parameters were calculated as: *α* = 4.43, *β* = 5.78. Data from *h ∈*0, 1 were combined, hence the mean value *h* = 0.5 was estimated.

**Figure S2.**
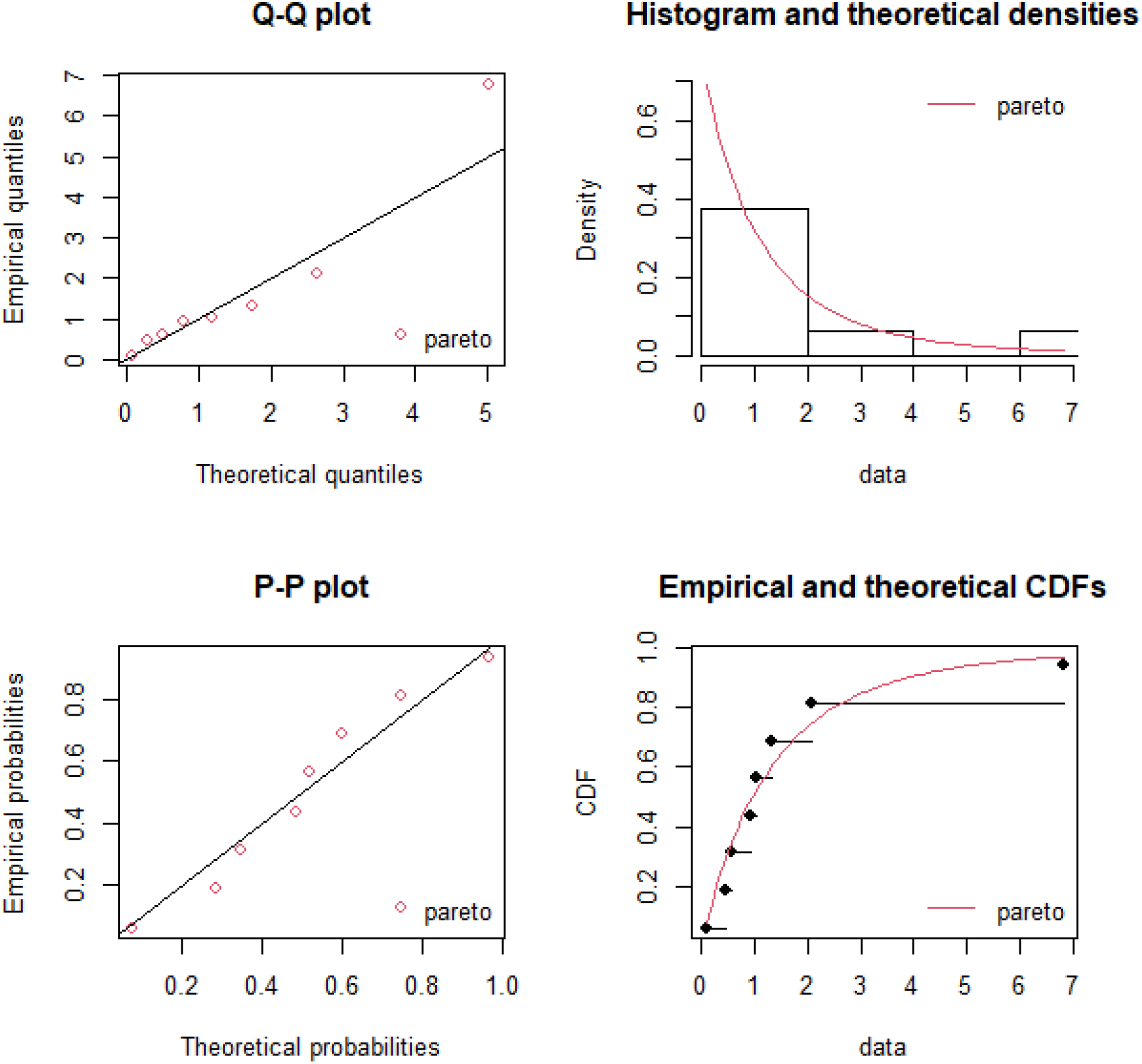
Pareto distribution analysis using ‘ fitdistrplus’, for DNA recovery following secondary transfer at times *T* |*h* = 0, 1, before rescaling with *k* = 0.67

For example, the probability, *Pr*(*S*|*h* = 0.5, *x* = 3) = 0.05 from eq S8. The corresponding value predicted by logistic regression of secondary transfer data is *Pr*(*S* |*h* = 0.5, *x* = 3) = 0.085 (fig. S3).

**Figure S3.**
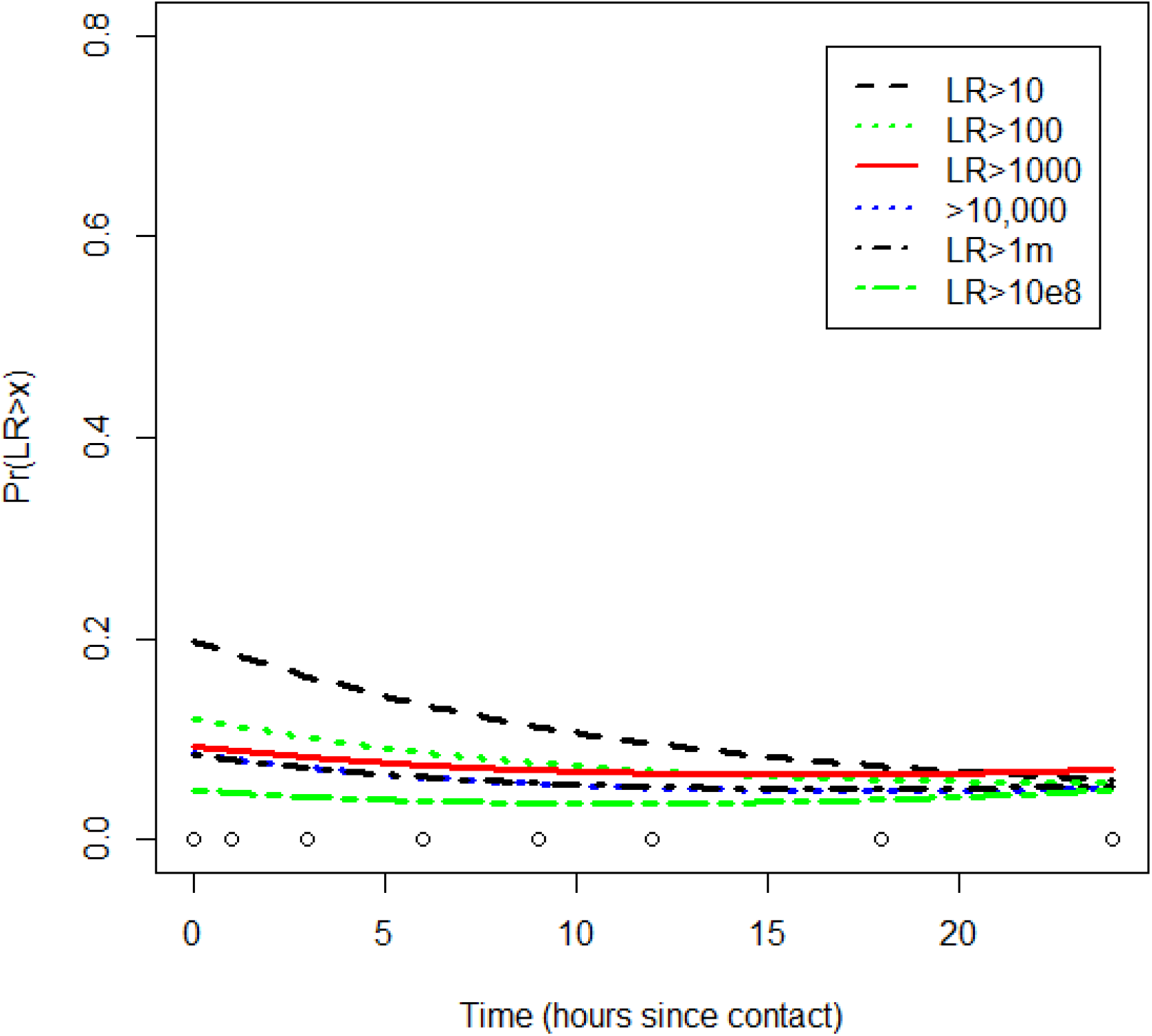
Secondary transfer logistic regression

The exponential distribution (*λ* = 0.598) returned similar results to Pareto (table S2).

**Table S2:** Comparison of Exponential vs. Pareto distributions of probability of secondary transfer (combination of h=0 and 1 data), relative to the logistic regression decision threshold (*x*).

| $Pr(log_{10}LR > x)$ | | |
| --- | --- | --- |
| $x$ | Exponential | Pareto |
| 1 | 0.183 | 0.163 |
| 2 | 0.101 | 0.088 |
| 3 | 0.055 | 0.052 |
| 4 | 0.030 | 0.032 |
| 6 | 0.009 | 0.014 |
| 8 | 0.003 | 0.007 |

### S4. Bayesian Logistic regression

The package rstanarm was used to prepare logistic regression models for a series of threshold values (*x* between 1 −10). Default priors chosen were weakly informative (Intercept and Time coefficients were modelled as: normal(location=0, scale (SD)=2.5)).

In R, the model is generated by the following command line:

~~~
Model=stan_glm∼Time,family=binomial(link=“logit”),dat)
~~~

Where ‘ dat’ is an array of the Time vs. *log*_10_*LR*_*ϕ*_ data for a given threshold value *x*.

For the output, 4000 randomly generated Intercept and Time coefficients were generated (Table S3). These coefficients are determined by MCMC; they define the shape of the logistic regression curve and are used to calculate probabilities of *s* or *t* conditioned on the time difference between sampling and deposition:

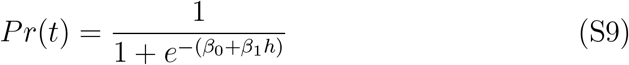

where *β*_0_ and *β*_1_ are the Intercept and Time coefficients respectively.

**Table S3:** Summary statistics of Direct transfer Bayesian logistic regression statistics generated from 4000 simulations per test. For each decision threshold *x*, there are two coefficients (Intercept and Time) that are used to generate the curves in 2. Summary statistics Rhat, n eff are discussed in S4.1. Quantiles for each coefficient are also listed

| Parameter | Rhat | n_eff | mean | sd | se_mean | 2.5% | 25% | 50% | 75% | 97.5% |
| --- | --- | --- | --- | --- | --- | --- | --- | --- | --- | --- |
| *log10LR > 1* |  |  |  |  |  |  |  |  |  |  |
| (Intercept) | 1.0000 | 2204 | 0.5762 | 0.3491 | 0.0074 | -0.0907 | 0.3386 | 0.5690 | 0.8061 | 1.2737 |
| Time | 1.0005 | 1707 | -0.1516 | 0.0448 | 0.0011 | -0.2457 | -0.1794 | -0.1490 | -0.1211 | -0.0695 |
| *log10LR > 2* |  |  |  |  |  |  |  |  |  |  |
| (Intercept) | 1.0009 | 2992 | 0.5091 | 0.3672 | 0.0067 | -0.2144 | 0.2672 | 0.5084 | 0.7445 | 1.2434 |
| Time | 1.0006 | 1946 | -0.1848 | 0.0493 | 0.0011 | -0.2878 | -0.2166 | -0.1823 | -0.1501 | -0.0946 |
| *log10LR > 3* |  |  |  |  |  |  |  |  |  |  |
| (Intercept) | 1.0006 | 2498 | 0.4469 | 0.3632 | 0.0073 | -0.2772 | 0.2043 | 0.4430 | 0.6913 | 1.1554 |
| Time | 1.0004 | 1654 | -0.2165 | 0.0583 | 0.0014 | -0.3413 | -0.2546 | -0.2120 | -0.1767 | -0.1102 |
| *log10LR > 4* |  |  |  |  |  |  |  |  |  |  |
| (Intercept) | 0.9996 | 2805 | 0.5164 | 0.3711 | 0.0070 | -0.2118 | 0.2674 | 0.5075 | 0.7484 | 1.2591 |
| Time | 1.0036 | 1341 | -0.2460 | 0.0645 | 0.0018 | -0.3863 | -0.2858 | -0.2431 | -0.1999 | -0.1327 |
| *log10LR > 5* |  |  |  |  |  |  |  |  |  |  |
| (Intercept) | 1.0013 | 2242 | 0.0846 | 0.3723 | 0.0079 | -0.6432 | -0.1647 | 0.0783 | 0.3249 | 0.8305 |
| Time | 1.0016 | 1323 | -0.2117 | 0.0641 | 0.0018 | -0.3455 | -0.2536 | -0.2084 | -0.1669 | -0.0982 |
| *log10LR > 6* |  |  |  |  |  |  |  |  |  |  |
| (Intercept) | 0.9998 | 2415 | -0.0500 | 0.3713 | 0.0076 | -0.7771 | -0.2984 | -0.0517 | 0.1995 | 0.6803 |
| Time | 1.0030 | 1528 | -0.1978 | 0.0592 | 0.0015 | -0.3233 | -0.2359 | -0.1951 | -0.1550 | -0.0934 |
| *log10LR > 8* |  |  |  |  |  |  |  |  |  |  |
| (Intercept) | 0.9998 | 2415 | -0.0500 | 0.3713 | 0.0076 | -0.7771 | -0.2984 | -0.0517 | 0.1995 | 0.6803 |
| Time | 1.0030 | 1528 | -0.1978 | 0.0592 | 0.0015 | -0.3233 | -0.2359 | -0.1951 | -0.1550 | -0.0934 |

These probabilities are input into the BN calculations described in section S2, using the logistic function, and the median value is calculated (quantiles can also be derived).

#### S4.1. Summary statistics

1. n eff is an estimate of the effective number of independent draws from the posterior distribution of the estimand of interest
2. Rhat: One way to monitor whether a chain has converged to the equilibrium distribution is to compare its behavior to other randomly initialized chains. This is the motivation for the Gelman and Rubin potential scale reduction statistic 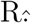 The Rŝtatistic measures the ratio of the average variance of samples within each chain to the variance of the pooled samples across chains; if all chains are at equilibrium, these will be the same and Rŵill be one. If the chains have not converged to a common distribution, the Rŝtatistic will be greater than one.
3. se mean: The standard error of the mean of the posterior draws (not to be confused with the standard deviation of the posterior draws) is the uncertainty associated with the Monte Carlo approximation. This quantity approaches 0 as the sample size goes to infinity.
4. sd: The posterior standard deviation of the posterior draws
5. The 2.5, 50 and 97.5 percentiles are shown

### S5. Calculation of the offset value

The Pareto distribution defines probability of secondary transfer eq S8:

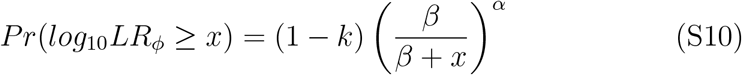

The probability of direct transfer is defined by the logistic regression in eq S9:

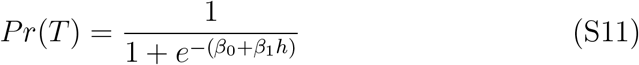

where *β*_0_ and *β*_1_ are the Intercept and Time coefficients respectively.

We seek to find (from eq: 5):

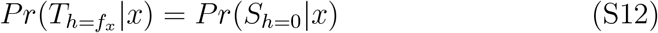

Where *f*_*x*_ is the offset value, which we need to determine from the following expression:

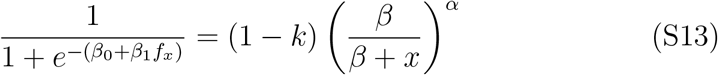

By rearrangement, to calculate *f*_*x*_:

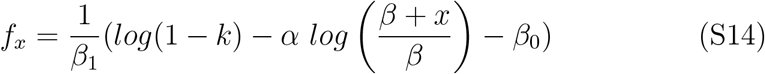

This expression is encoded into ALTRaP and is used to determine the offset value for each of the 4000 simulations per value of *x*.

Note that for the *Pr*(*S*) sensitivity tab in ALTRaP, the value is determined by optimisation.

### S6. The case of Queen v. William Francis Jones

From the court of England and Wales, Queen v. William Francis Jones, the defendant’ s conviction was quashed [33]. In this case a DNA profile was found on the pin of a hand-grenade and a statement was provided at trial:

At the original trial this evidence was sufficient for conviction.

At appeal, the defence accepted the prosecution’ s sub-source proposition:

*“that there could not be a realistic challenge to the inference that the DNA sample obtained from the firing pin came from his client”*.

The next step was to consider the evidence at activity level, noting that this aspect had not been adequately dealt with at the original trial.

The court of appeal directed two experts representing the prosecution and defence to prepare a joint statement (for details of the rules see the Forensic Science Regulator Codes of Practice and Conduct [32]). It is important the courts provide a framework for experts to be able to debate and to be able to agree their common ground outside the court-room. The procedure becomes *collaborative*, rather than *combative*. Once the joint report is written, the areas of agreement are clear and areas of disagreement can be explored further if required.

In this case, the experts had no disagreements and wrote:

*“The statistical evaluation provided addresses only whether an individual could be a possible donor of DNA and does not address the mechanism by which any DNA was deposited, the time at which it may have been deposited, or the order in which different contributions of DNA were deposited*.….

*If it were to be accepted that the DNA from William Jones is present on the safety pin, then the DNA result alone does not assist in determining: (a) whether William Jones was the last person to touch the safety pin before it was recovered; (b) how long ago the DNA from William Jones was deposited on the safety pin; (c) the mechanism by which the DNA from William Jones was deposited on the safety pin, including whether it was left directly (primary transfer) or indirectly via an intermediary (secondary transfer)”*

On the basis of the expert’ s report the conviction was quashed. In the absence of data or a model to assist the analysis of the evidence at activity level, the scientists were unable to help the court with further with their deliberations. The limitations of evidence at sub-source level were made clear. Consequently, the conviction was quashed.

This case is a vivid reminder of the dangers of not alerting the court to the limitations of sub-source level reporting. Lack of education of experts, judges and lawyers, along with an unfavourable ‘ combative’ court environment are all barriers to proper exposition of the evidence.

## References

[1] P. Gill, T. Hicks, J. M. Butler, E. Connolly, L. Gusmão, B. Kokshoorn, N. Morling, R. A. van Oorschot, W. Parson, M. Prinz, et al., DNA commission of the international society for forensic genetics: Assessing the value of forensic biological evidence-guidelines highlighting the importance of propositions. Part II: Evaluation of biological traces considering activity level propositions, Forensic Science International: Genetics 44 (2020) 102186.

[2] Z. E. Bowman, K. S. Mosse, A. M. Sungaila, R. A. van Oorschot, D. Hartman, Detection of offender DNA following skin-to-skin contact with a victim, Forensic science international: genetics 37 (2018) 252– 259.

[3] D. Taylor, A. Biedermann, T. Hicks, C. Champod, A template for constructing bayesian networks in forensic biology cases when considering activity level propositions, Forensic Science International: Genetics 33 (2018) 136–146.

[4] J. M. Butler, Forensic DNA typing: biology, technology, and genetics of STR markers, Elsevier, 2005.

[5] Ø. Bleka, G. Storvik, P. Gill, Euroformix: an open source software based on a continuous model to evaluate STR DNA profiles from a mixture of contributors with artefacts, Forensic Science International: Genetics 21 (2016) 35–44.

[6] D. Taylor, J.-A. Bright, C. McGoven, C. Hefford, T. Kalafut, J. Buckleton, Validating multiplexes for use in conjunction with modern interpretation strategies, Forensic Science International: Genetics 20 (2016) 6–19.

[7] O. Hansson, P. Gill, Characterisation of artefacts and drop-in events using STR-validator and single-cell analysis, Forensic Science International: Genetics 30 (2017) 57–65.

[8] C. Muth, Z. Oravecz, J. Gabry, User-friendly bayesian regression modeling: A tutorial with rstanarm and shinystan, Quantitative Methods for Psychology 14 (2) (2018) 99–119.

[9] M. Goray, E. Eken, R. J. Mitchell, R. A. van Oorschot, Secondary DNA transfer of biological substances under varying test conditions, Forensic Science International: Genetics 4 (2) (2010) 62–67.

[10] M. L. Delignette-Muller, C. Dutang, et al., fitdistrplus: An R package for fitting distributions, Journal of statistical software 64 (4) (2015) 1– 34.

[11] E. A. Dowlman, N. C. Martin, M. J. Foy, T. Lochner, T. Neocleous, The prevalence of mixed DNA profiles on fingernail swabs, Science & Justice 50 (2) (2010) 64–71.

[12] S. Willis, L. McKenna, S. McDermott, G. O’Donell, A. Barrett, B. Rasmusson, A. Nordgaard, C. Berger, M. Sjerps, J. Lucena-Molina, et al., Enfsi guideline for evaluative reporting in forensic science, European Network of Forensic Science Institutes. !http://enfsi.eu/wp-content/uploads/2016/09/m1_guideline.pdf (2015).

[13] D. Taylor, T. Hicks, C. Champod, Using sensitivity analyses in bayesian networks to highlight the impact of data paucity and direct future analyses: a contribution to the debate on measuring and reporting the precision of likelihood ratios, Science & Justice 56 (5) (2016) 402–410.

[14] S. Zoppis, B. Muciaccia, A. D’Alessio, E. Ziparo, C. Vecchiotti, A. Filippini, DNA fingerprinting secondary transfer from different skin areas: morphological and genetic studies, Forensic Science International: Genetics 11 (2014) 137–143.

[15] D. Taylor, B. Kokshoorn, A. Biedermann, Evaluation of forensic genetics findings given activity level propositions: a review, Forensic Science International: Genetics 36 (2018) 34–49.

[16] R. A. van Oorschot, B. Szkuta, G. E. Meakin, B. Kokshoorn, M. Goray, DNA transfer in forensic science: a review, Forensic Science International: Genetics 38 (2019) 140–166.

[17] M. Goray, R. A. van Oorschot, The complexities of DNA transfer during a social setting, Legal Medicine 17 (2) (2015) 82–91.

[18] R. van Oorschot, D. McColl, J. Alderton, M. Harvey, R. Mitchell, B. Szkuta, Activities between activities of focus—relevant when assessing DNA transfer probabilities, Forensic Science International: Genetics Supplement Series 5 (2015) e75–e77.

[19] D. Taylor, A. Biedermann, L. Samie, K.-M. Pun, T. Hicks, C. Champod, Helping to distinguish primary from secondary transfer events for trace DNA, Forensic Science International: Genetics 28 (2017) 155–177.

[20] J. Helmus, T. Bajanowski, M. Poetsch, DNA transfer—a never ending story. a study on scenarios involving a second person as carrier, International journal of legal medicine 130 (1) (2016) 121–125.

[21] B. Szkuta, K. N. Ballantyne, B. Kokshoorn, R. A. van Oorschot, Transfer and persistence of non-self DNA on hands over time: using empirical data to evaluate DNA evidence given activity level propositions, Forensic Science International: Genetics 33 (2018) 84–97.

[22] M. van den Berge, G. Ozcanhan, S. Zijlstra, A. Lindenbergh, T. Sijen, Prevalence of human cell material: DNA and RNA profiling of public and private objects and after activity scenarios, Forensic Science International: Genetics 21 (2016) 81–89.

[23] M. Bouzga, G. Dørum, K. Gundersen, P. Kohler, P. Hoff-Olsen, A. Fonneløp, Is it possible to predict the origin of epithelial cells?–a comparison of secondary transfer of skin epithelial cells versus vaginal mucous membrane cells by direct contact, Science & Justice (2020).

[24] C. Ladd, M. S. Adamowicz, M. T. Bourke, C. A. Scherczinger, H. C. Lee, A systematic analysis of secondary DNA transfer, Journal of Forensic Science 44 (6) (1999) 1270–1272.

[25] C. Champod, DNA transfer: informed judgment or mere guesswork?, Frontiers in genetics 4 (2013) 300.

[26] P. Gill, T. Hicks, J. M. Butler, E. Connolly, L. Gusmão, B. Kokshoorn, N. Morling, R. A. van Oorschot, W. Parson, M. Prinz, et al., DNA commission of the international society for forensic genetics: Assessing the value of forensic biological evidence-guidelines highlighting the imporance of propositions: Part I: evaluation of DNA profiling comparisons given (sub-) source propositions, Forensic Science International: Genetics 36 (2018) 189–202.

[27] L. Samie, C. Champod, D. Taylor, F. Taroni, The use of bayesian networks and simulation methods to identify the variables impacting the value of evidence assessed under activity level propositions in stabbing cases, Forensic Science International: Genetics (2020) 102334.

[28] B. Efron, R. J. Tibshirani, An introduction to the bootstrap, CRC press, 1994.

[29] E. A. M. Graham, G. N. Rutty, Investigation into “normal” background DNA on adult necks: implications for DNA profiling of manual strangulation victims, Journal of forensic sciences 53 (5) (2008) 1074–1082.

[30] E. A. M. Graham, W. J. Watkins, F. Dunstan, S. Maguire, D. Nuttall, C. E. Swinfield, G. N. Rutty, A. M. Kemp, Defining background DNA levels found on the skin of children aged 0–5 years, International journal of legal medicine 128 (2) (2014) 251–258.

[31] R. v Peter Weller [2009] EWCA Crim 1670. Retrieved from: https://www.bailii.org/ew/cases/EWCA/Crim/2010/1085.html (2009).

[32] Forensic Science Regulator Codes of Practice and Conduct: Development of Evaluative Opinions FSR-C-118 Issue 1. Retrieved from:https://assets.publishing.service.gov.uk/government/uploads/system/uploads/attachment_data/file/960051/FSR-C-118_Interpretation_Appendix_Issue_1002_.pdf (2021).

[33] Jones v R. [2020] EWCA Crim 1021. Retrieved from: https://www.bailii.org/ew/cases/EWCA/Crim/2020/1021.html (2020).

[34] P. Gill, Interpretation continues to be the main weakness in criminal justice systems: Developing roles of the expert witness and court, Wiley Interdisciplinary Reviews: Forensic Science 1 (2) (2019) e1321.

[35] F. Taroni, S. Bozza, A. Biedermann, C. Aitken, Dismissal of the illusion of uncertainty in the assessment of a likelihood ratio, Law, Probability and Risk 15 (1) (2016) 1–16.

[36] A. Biedermann, S. Bozza, F. Taroni, C. Aitken, The meaning of justified subjectivism and its role in the reconciliation of recent disagreements over forensic probabilism, Science & Justice 57 (6) (2017) 477–483.

[37] G. E. Meakin, A. Jamieson, A response to a response to meakin and jamieson DNA transfer: review and implications for casework, Forensic Science International: Genetics 22 (2016) e5–e6.

[38] L. Samie, F. Taroni, C. Champod, Estimating the quantity of transferred DNA in primary and secondary transfers., Science & justice: journal of the Forensic Science Society 60 (2) (2020) 128.

[39] S. Højsgaard, D. Edwards, S. Lauritzen, Graphical models with R, Springer Science & Business Media, 2012.

